# Modelling mechanochemical coupling in optogenetically activated cell layers

**DOI:** 10.1101/2025.06.30.662367

**Authors:** Dennis Wörthmüller, Falko Ziebert, Ulrich S. Schwarz

## Abstract

In adherent cells, actomyosin contractility is regulated mainly by the RhoA signaling pathway, which can be controlled by optogenetics. To model the mechanochemical coupling in such systems, we introduce a finite element framework based on the discontinuous Galerkin method, which allows us to treat cell doublets, chains of cells and monolayers within the same conceptual framework. While the adherent cell layer is modeled as an actively contracting viscoelastic material on an elastic foundation, different models are considered for the Rho-pathway, starting with a simple linear chain that can be solved analytically and later including direct feedback that can be solved only numerically. Our model predicts signal propagation as a function of coupling strength and viscoelastic time scales and identifies the conditions for optimal cell responses and wave propagation. In general, it provides a systematic understanding of how biochemistry and mechanics simultaneously contribute to the communication of adherent cells.

## Introduction

With the advent of mechanobiology, it has become clear that biochemistry and mechanics play an equally important role for the function of cellular systems (1, 2). For example, it has been shown that stem cells actively sense the stiffness of their environment, and that this mechanical input together with soluble factors determines their subsequent differentiation (3, 4). Later it has been shown that one important element is the transcription factor Yes-associated protein (YAP), which is activated on stiff substrates and in geometrical confinement (5, 6).

A close coupling between biochemistry and mechanics also exists for multicellular systems. For example, it has been shown that in wound healing and infection, neighboring epithelial cells coordinate their activity by mechanical activation of extracellular signal-regulated kinases (ERK) (7, 8). A mathematical model demonstrated how wave propagation results from mutual feedback between mechanics and biochemistry: leader cells pull on their followers, ERK is activated in the followers and activates contractility, which leads to forces on the next row of followers (9–11). A similar mechanism is also realized by the tumour suppressor protein merlin, which under mechanical force is relocalized from the cell-cell junctions to the cytoplasm (12). Interestingly, the mechanism for wave propagation is similar to the one for action potentials in the neurosciences. In principle, the signal could go both ways, but a refractory period in the sending part prevents the wave from going backwards.

The general scheme of mechanochemical feedback leading to complex systems behaviour is even more evident for developing organisms (13, 14). Here morphogen concentration fields determine where cells grow and divide, and this leads to changes of the domain in which signaling is active, which in turn changes the way morphogens are secreted and distributed. Such feedback loops lead to non-linear systems dynamics which can explain the intricate patterns that emerge during tissue formation and embryogenesis (13). One prominent model organism is the fruit fly Drosophila, where experimental observations have been coupled with mathematical models (15, 16). Another one is the freshwater polyp Hydra, which is able to regenerate its patterning even after being cut in pieces (17–20). However, for such organismal systems it is very challenging to achieve a systems level understanding connecting molecular processes to the tissue scale. For mechanistic understanding, it is therefore rewarding to turn back to their elementary building blocks, the cells, and small assemblies of such cells, with the long term aim to upscale to larger systems.

To understand the coupling of biochemistry and mechanics at the level of single cells, one must start with the actin cytoskeleton (21–23). This is a network of actin filaments and myosin II molecular motor proteins that determine the mechanical properties of cells, particularly during processes such as adhesion, migration, and division. Since the actin cytoskeleton continuously consumes energy in the form of adenosine triphosphate (ATP) to grow and reorganize its filaments, generating forces and flows in the process, the appropriate modeling approach is to introduce active stresses that are coupled to the chemical potential of the myosin II motors. This central concept led to the development of active gel theory (24, 25). Active gel theory is commonly used to model single-cell migration (26, 27), but can also be readily extended to larger systems, such as tissue flow (28).

To incorporate more mechanistic details into the process of force generation by cells, one must also consider how it is regulated by the small GTPases from the Rho-family, including RhoA, Rac1 and Cdc42 (23, 29). Briefly, Rac1 and Cdc42 primarily regulate the assembly of larger protrusive actin structures, such as lamellipodia and filopodia, respectively, while RhoA is predominantly responsible for the formation of actomyosin contractility. The activity of these small GTPases is controlled by many different Guanine Exchange Factors (GEFs). If such a GEF activates RhoA, this in turn activates Diaphanous-related formin (Dia) for actin polymerization and Rho-associated protein kinase (ROCK) for contractility through myosin II molecular motors (30). Together, these effects then lead to productive force generation.

Several positive and negative feedback loops exist among these components, leading to complex temporal dynamics and pattern formation (23, 29). For instance, Bement et al. (31) identified an activator-inhibitor relationship between RhoA and F-actin, which results in the emergence of spiral contraction waves during cytokinesis in Xenopus embryonic cells. Similar surface contraction waves have been observed in starfish oocytes during maturation, modeled by the coupled reaction kinetics of actin and myosin II (32, 33). In a systematic study, by combining nonlinear system dynamics and experimental data, Kamps et al. (34) developed a detailed model for the reaction kinetics of GEF, RhoA, and myosin This model not only demonstrates the complexity of the RhoA pathway, but also successfully explains the experimental observations of pulsatile contractions in the actin cortex, identifying cytosolic GEFH1 as a crucial parameter for the emergence of this pulsatile behavior.

While GEFs for the small GTPases from the Rho-family sometimes are activated purely by biochemical pathways, often their activation results from mechanical forces (35–37), similar to the cases of ERK (7, 9, 10) and merlin (12). Often, theoretical models incorporate these concepts by proposing feedback mechanisms between mechanical tension and biochemical signaling. For example, a positive biochemical-mechanical feedback loop between forces exerted on focal adhesions and RhoA signaling at these sites can explain spatial gradients in the periodic myosin-*α*-actinin pattern in stress fibers stimulated with calyculin A (38). Several mathematical studies combined simple models for Rho GTPase activity and cell mechanics to demonstrate that their interplay leads to complex cell behaviors (39, 40). The authors showed that their proposed system can exhibit bistability, where the two states represent permanently contracted or relaxed cells, and can also produce oscillatory states. Recently, Staddon et al. (41) coupled a basic activator-inhibitor reaction-diffusion (RD) system, comprising RhoA as the activator and myosin II as the inhibitor, with the mechanics of viscoelastic solids and fluids. In this model, the interplay between biochemistry, actomyosin contractility, and viscoelastic deformation leads to the emergence of propagating pulsatile contractions and topological turbulence in flows of RhoA.

In order to dissect these signaling pathways experimentally, one usually works with inhibitors, which are small chemical molecules that reduce the effect of certain components like ROCK or myosin II. However, the concomitant results are often rather qualitative in nature and it is not always clear how well the inhibitor reaches its putative target. Recently, optogenetics has emerged as a powerful alternative, which leads to more quantitative results (42). In optogenetics, a light-sensitive construct is engineered into the cells and can then be activated with high temporal and spatial resolution. Among others, this approach has been applied in several studies for example to control neural activity (43), the regulation of gene expression (44, 45) or even to regulate engineered metabolic pathways in cells (46), which illustrates the versatility of this method. In the field of mechanobiology it has become an established technique to activate the Rho-pathway by recruiting a GEF to the membrane (33, 47–49), thus allowing for a precise spatiotemporal control of cytoskeletal dynamics of single cells (50, 51) and multicellular systems (52, 53).

The quantitative advances achieved by optogenetics now open the door for a more detailed mathematical modelling of the underlying processes. Different modelling frameworks have been applied before to couple biochemistry and mechanics. One attractive option is the cellular Potts model, which has been applied to both Rho/Rac-signaling (54) and EKR-signaling (7). However, the framework is not able to model active stresses in detail. Here we therefore turn to active continuum mechanics, which is the natural framework to describe local active stresses, as demonstrated by the success of active gel theory (24–28). For the signaling pathways, we rely on a description in terms of a system of RDequations (34) solved on a geometrical domain which represents a cell ensemble strongly adhered to an elastic foundation (50, 55, 56). Because both frameworks, continuum mechanics and RD-systems, lead to partial differential equations (PDEs), we turn to the finite element method (FEM), which is a standard way to numerically solve PDEs. In particular, since we aim to model multicellular systems, for the RD-part we implement the discontinuous Galerkin (DG) version of FEM, which offers a natural way to represent the discontinuities in concentrations at the cell-cell boundaries in multicellular systems. Here we first lay the conceptual basis for such an approach and then present representative applications for mechanochemical pattern formation in multicellular systems controlled by optogenetics.

The manuscript is structured as follows. First we will introduce the main concepts and equations, motivated by recent experiments on optogenetic activation of cell doublets (52). Our starting point is the observation that optogenetic activation of contractility in one cell triggers an active response in a neighboring cell, compare Fig. 1a. For Rho-activation, we start with a simple linear variant of the pathway, which is sufficient to describe the recent experiments. We then explore the consequences of this response, going from the cell doublet on a H-pattern to increasingly larger systems, namely cell chains and monolayers as commonly used in experiments (52, 53), compare Fig. 1b. Finally, we will demonstrate the generality of the simulation framework by addressing the case of a monolayer with a more dynamic model for the reaction kinetics of the Rho-pathway (34).

**Fig. 1.**
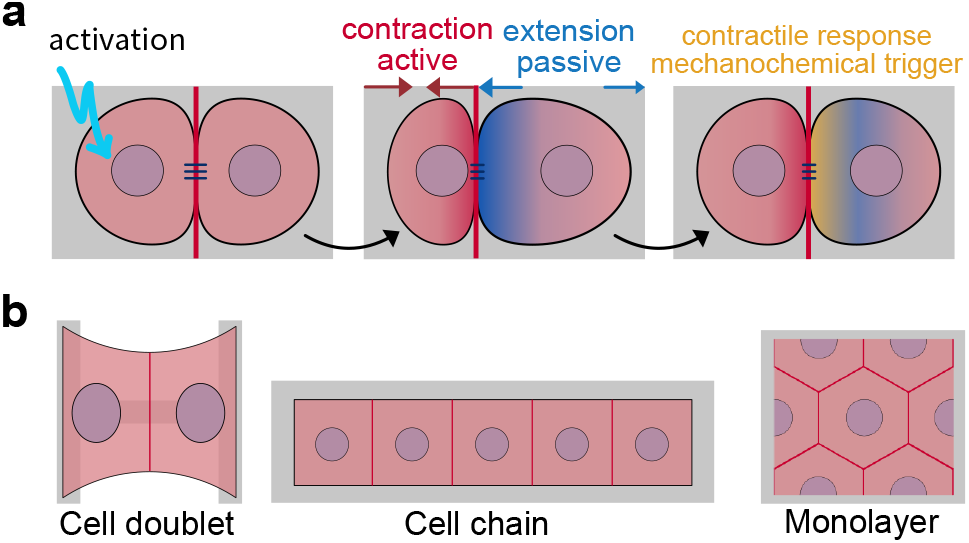
Mechanochemical coupling in different geometries. Cells in contact with each other communicate through mechanochemical coupling. In the cell doublet, the optogenetically controlled contraction of the left cell induces an active contractile response in the right cell. (b) For spatial modelling, we specify adhesion geometries commonly used in experiments. We start with the cell doublet on an H-pattern and then continue to cell chains and cell monolayers.

## Model

### Coupling biochemistry and mechanics

Our modelling approach is strongly motivated by recent experiments on cell doublets on a H-pattern whose contractility is activated in the left cell by Rho-optogenetics, such that one can follow the response of the right cell in quantitative detail (52). The formulation of the model follows the central observation that actively generated stresses within the cell layer depend on the concentration of the downstream output of the RhoA-pathway. Assuming that sufficient amounts of actin filaments are generated by the Dia-leg, this is mainly the amount of active myosin II generated by the ROCK-leg of the pathway. The spatio-temporal distribution of actively generated stresses depends also on the reaction kinetics and diffusive properties of all upstream signalling proteins. The active stresses may then lead to deformation of the cell which directly feeds back to the RD-system by generating advection terms and changing concentrations. Consequently, the spatiotemporal evolution of a signaling protein concentration *c*_*i*_(**x**, *t*) is described by a reaction-diffusion-advection equation on a two-dimensional time-dependent domain Ω(*t*),

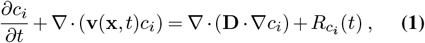

where **D** denotes the two-dimensional diffusion tensor, *R*_*ci*_ the reaction kinetics and the index *i* represents a signaling protein in the RhoA pathway. Eq. (1) arises naturally by demanding local mass conservation on the time-dependent domain by following Reynold’s transport theorem. It includes an advection term **v ∇***c*_*i*_ due to flows induced by contraction and expansion and an enrichment/dilution term *c*_*i*_ ∇ **v** due to local volume changes, where **v**(**x**, *t*) corresponds to the velocity of the deforming material. Hence, deformations naturally interfere with the spatiotemporal evolution of the protein concentrations. Eq. (1) is written in terms of spatial (Eulerian) coordinates **x** which is a convenient choice for the description of diffusion processes.

However, the deformation of the cell domain is better treated in terms of referential (Lagrangian) coordinates 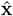. The two coordinate systems are related by the deformation field 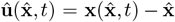. We use Piola’s identity 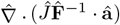, where **a** is an arbitrary vector field and 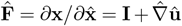 the deformation gradient tensor with 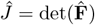, to pull Eq. (1) back to the reference configuration Ω_0_ and express it in terms of Lagrangian coordinates as

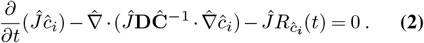

Here, 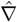 denotes the derivative with respect to Lagrangian coordinates, **D** = *D***I** is assumed to be an isotropic tensor with scalar diffusivity *D* and 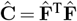 denotes the Cauchy-Green deformation tensor. The velocity of the material is absorbed into the time-derivative in the first term. For the details of this derivation we refer to the supporting information. From the first term in Eq. (2) we see that compression (∂_*t*_*Ĵ <* 0) and dilation (∂_*t*_*Ĵ >* 0) of the elastic domain effectively alters the reaction kinetics. Further, we see that the diffusion is impacted by local deformations and can become anisotropic. Following earlier work on cells as active materials, we describe the mechanics of the cell layer as a viscoelastic continuum with active stresses coupled to an elastic foundation (50, 55, 57, 58) (compare Fig. 2a). In particular, we assume that the lateral extent of the cell layer is much larger than the thickness *h*_*c*_ of its effective contractile layer at the basal side. This assumption also underlies the widely used method of monolayer stress microscopy (52, 59–61). Note that *h*_*c*_ is usually not larger than one micrometer, while the cell height for strongly adhering cells typically varies between two and ten micrometers, depending on cell type, cell density and shape of the nuclei. The thin layer assumption allows us to obtain a two-dimensional model by using the plane stress conditions for the elastic sheet. The force balance equation then reads

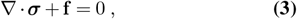

where ***σ*** is the two-dimensional in-plane Cauchy stress tensor and **f** denotes an externally applied two-dimensional body force. The two-dimensional stress tensor is obtained by averaging the three-dimensional stress tensor over the thickness of the cell layer.

**Fig. 2.**
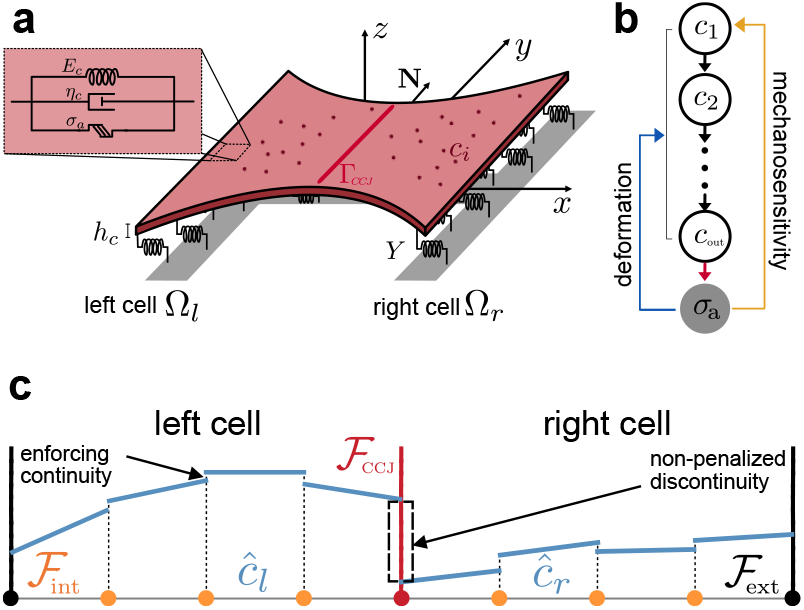
Mechanochemical model for a cell doublet. (a) A cell doublet (Young’s modulus *E*_*c*_, viscosity *η*_*c*_, active stress *σ*_*a*_, contractile layer thickness *h*_*c*_) adheres to an elastic foundation via springs with stiffness density *Y* that are homogeneously distributed over the area defined by the H-shaped micropattern. The cell-cell junction (red line; Γ_CCJ_) separates the two cells (Ω_*l*_, Ω_*r*_) along the vertical symmetry axis of the micropattern (y-axis). (b) The simplest model for the signaling cascade is a linear chain in which mechanosensitivity leads to activation of the first signaling protein and the last signaling protein determines active stresses. (c) Discontinuous Galerkin method for the reaction-diffusion system. Solutions are naturally allowed to have discontinuities across the mesh facets. Penalizing discontinuities at internal facets ℱ_int_ (orange dots) leads to approximately smooth solutions within cells. Dropping this penalty at the cell-cell boundary ℱ_CCJ_ results in jumps.

For the cell layer, we assume a linear viscoelastic constitutive relation of the solid (Kelvin-Voigt) type. Although cells are very dynamic and often are modelled by viscoelasticity of the fluid (Maxwell) type, especially in active gel theory (25), here we consider stably adhering cells that effectively behave as solids due to homeostatic mechanisms, including volume control. The Kelvin-Voigt law reads

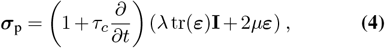

where ***ε*** = (∇**u** + ∇**u**^T^)*/*2 denotes the infinitesimal strain tensor and *τ*_*c*_ = *η*_*c*_*/E*_*c*_ is the relaxation time defined by the ratio of viscosity *η*_*c*_ and Young’s modulus *E*_*c*_ of the cell. Further, we introduce the two-dimensional Lamé coefficients as

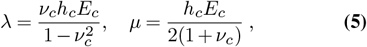

where *ν*_*c*_ is the cellular Poisson’s ratio.

Actomyosin contractility is modeled by an active stress tensor ***σ***_a_ and the total cell stress is given by the sum of the passive and active contributions, ***σ*** = ***σ***_p_ + ***σ***_a_. The cell layer is coupled to an elastic substrate which can be thought of as a continuous layer of springs between the cell and a rigid substrate (55, 62). In particular, the elastic foundation can also describe an elastic substrate as commonly used in traction force microscopy. It is described by a local force per unit area

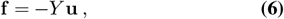

where *Y* is the spring constant density ([*Y*] = Nm^−3^) and **u** the displacement field of the cell layer. In case of a micropatterned surface, the spring constant density becomes position dependent and defines the adhesion geometry. Note that ***σ*** measures stresses in the deformed configuration and is evaluated at spatial coordinates. This means that in order to express Eq. (3) in terms of referential coordinates one has to use the first Piola-Kirchhoff stress tensor 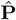. This distinction becomes important only in the limit of finite strains. In the limit of small strains, i.e. small deformation gradients, we can assume 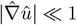 and only consider terms up to linear order in 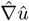 for which we can approximate 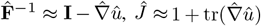 and 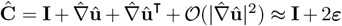 Given this small strain assumption, the two tensors only differ in terms 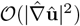, such that 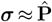. We will use referential coordinates for the force balance and constitutive relation, respectively.

The boundary of the cells is non-permeable for the signaling proteins and hence, we impose zero-flux boundary conditions at the interface between cell interior and cell exterior **j·N** = 0 on ∂Ω_0_ with 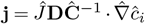 being the diffusive flux. Besides ensuring mechanical integrity, an inherent feature of intercellular junctions, e.g. tight junctions, is to maintain compartmentalisation in tissues by acting as a barrier for fluids and solutes. Therefore also the cell-cell junctions (CCJ) are non-permeable, which is incorporated by imposing an internal zero-flux boundary condition on Γ_CCJ_. As no external stresses are applied at the boundary ∂Ω_0_ of the sheet, we have the boundary condition ***σ*** · **N** = 0 on ∂Ω_0_.The net traction force exerted by the cell layer vanishes as 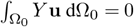, as required for a closed system.

In summary, the model can be formulated as follows: Find the displacement field **u** together with the concentrations *ĉ*_*i*_ of the signaling species such that

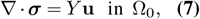

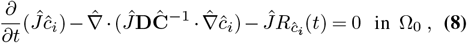

together with the boundary conditions

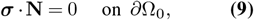

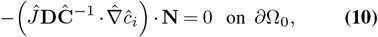

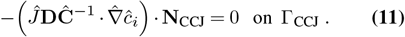

In addition to the geometrically arising coupling between the RD-system and mechanics as described by Eqs. (7, 8), we introduce two additional coupling mechanisms as schematically illustrated in Fig. 2b. First, we couple the output of the RD-system, i.e. the concentration of the last signaling protein of the activation cascade *c*_out_, to active stresses *σ*_a_ (red arrow in Fig. 2b). For this we relate 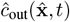 to *σ*_a_ via a relation (41, 63)

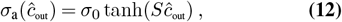

where *σ*_0_ is the maximal contractile stress and *S* a parameter which controls how sensitive stress generation is. Since tanh(*x*) ≤ 1 everywhere, we additionally ensure that active stresses are bound and hence avoid numerical instabilities. Second, we introduce mechanosensitivity by relating the mechanical perturbations to the activation of the most upstream signaling protein *ĉ*_1_ in the RD-system, see Fig. 2b (yellow arrow). In our model, the mechanical perturbation can either be force-related (measured in terms of internal stresses) or deformation-related (expressed in terms of strain or compression/stretch). From the perspective of continuum mechanics, these measures are provided by the Cauchy stress tensor ***σ*** and the Cauchy strain tensor ***ε*** (or the deformation gradient tensor **F**). To make this coupling independent of the frame of reference, for each of the different measures we can choose between two tensor invariants (for a 2D system), the trace or the determinant (64). Motivated by experimental studies in a strain-controlled experimental setup, showing that cells respond directly to stretch (65), we decide to introduce a strain-dependent feedback. This choice implies that we use the trace of the strain tensor, tr(***ε***), because the determinant det(***ε***) is at least of order *ε*^2^ and not suitable for a linear constitutive relation. However, in more detailed models of mechanotransduction, higher order invariants could be included. In this case one also had to go to non-linear material laws, e.g. Neo-Hookean, for consistency. We also note that mechanosensitive proteins usually respond to extension, not to compression, and thus we use strain only if it is extensile. Hence, we assume an activation rate of the first signaling protein due to passive strains via a source term of the form

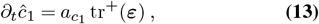

where *a*_*c*1_ is a generic activation rate which might depend on other quantities depending on the specific choice for the RD-system (similar to Hino et al. (7)). The plus sign indicates that coupling is only present in regions of positive strains, i.e. tr^+^(***ε***):= max (0, tr(***ε***)). We want to emphasize that the magnitude of *a*_*c*1_ has to be chosen such that if multiplied by tr^+^(***ε***), it results in an appropriate rate for the strain-dependent feedback.

### Discontinuous Galerkin method for reaction-diffusion system

To capture sharp concentration changes at cell-cell junctions in our RD-system. we employ a discontinuous Galerkin (DG) finite element scheme to discretize the pulled-back Eq. (8). If a standard continuous Galerkin (CG) formulation were used, the concentration field *ĉ* would be forced to remain continuous across all element boundaries (mesh facets ℱ), including the element boundaries representing the CCJ. However, the relevant signaling proteins do not cross cell-cell boundaries, meaning concentrations may differ significantly between neighboring cells and fluxes across the cell-cell boundary remain zero. Using CG, this would require solving a separate system of partial differential equations in each cell and manually enforcing the interface conditions. By contrast, DG is a natural choice for concentration differences at cell-cell interfaces. The key idea of this method is sketched in Fig. 2c. In DG the basis functions are not required to be continuous across facets. Continuity can instead be enforced in a weak sense by adding a penalty term that discourages jumps of the solution across facets. On internal facets ℱ_int_ (orange dots in Fig. 2c), we apply this penalty to obtain an approximately continuous solution within each cell (small jumps at facets). On facets ℱ_CCJ_ representing the CCJs (red dots in Fig. 2c), however, we relax this penalization, naturally allowing concentrations to vary between cells (large jumps at facets) while ensuring that no flux crosses the CCJ. Zero flux across external facets ℱ_ext_ is ensured by natural boundary conditions Eq. (10). In practice we use the symmetric weighted interior penalty (SWIP) variant (66), which is particularly well suited for problems with heterogeneous diffusion (Eq. (8)). A derivation of the weak form is provided in the supplemental text. The mechanical part of our model is solved with standard CG, which is sufficient because displacements are continuous across the cell-cell boundary.

### Simple model for the Rho-pathway with optogenetic activation

Although the RhoA-pathway allows for complex dynamics, earlier work has shown that at least in cells with strong adhesion, the actin cytoskeleton is regulated in the vicinity of a stable fixed point of this pathway (50, 52). One aspect could be that strongly adherent cells exhibit dominant stress fibers. Since stress fibers are highly organized structures, it is plausible to assume a differently organized RD-system than for e.g. the homogeneous actin cortex in egg cells (31). Indeed it has been shown recently that different organizations of the actin cytoskeleton lead to different activation and relaxation times in the Rho-pathway (50). Another aspect might be the observation that the stability of the RhoA-pathway has a strong dependence on the total GEF-concentration (34). This suggests that the total GEF expression levels, which are naturally elevated in cells transfected with an optogentic construct, render their RhoA-system more stable. Since cells transfected with the CRY2/CIBN system show a significantly higher baseline contractility, we assume that this might correspond to the stable branch of high GEF-concentrations (34).

First, we want to focus on this regime and describe optogenetic activation as a reversible process such that after activation cells eventually go back to their homeostatic contractility level without showing any significant oscillatory or excitable behavior upon photoactivation. Excitability is therefore completely controlled by the recruitment of a GEF to the membrane and can be scaled by the duration of the activation light pulse until saturation sets in (47, 48). Motivated by these observations, we first assume a linear input-output relationship between GEF plasma membrane recruitment and myosin II induced contractility. This assumption not only reduces the number of unknown parameters, it also allows for an analytical solution of the homogeneous system, such that it can be fully understood. Our proposed RhoA-myosin reaction scheme is shown in Fig. 3a. GEF activity enters implicitly through a predefined input signal which we describe by a function

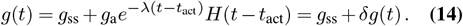

**Fig. 3.**
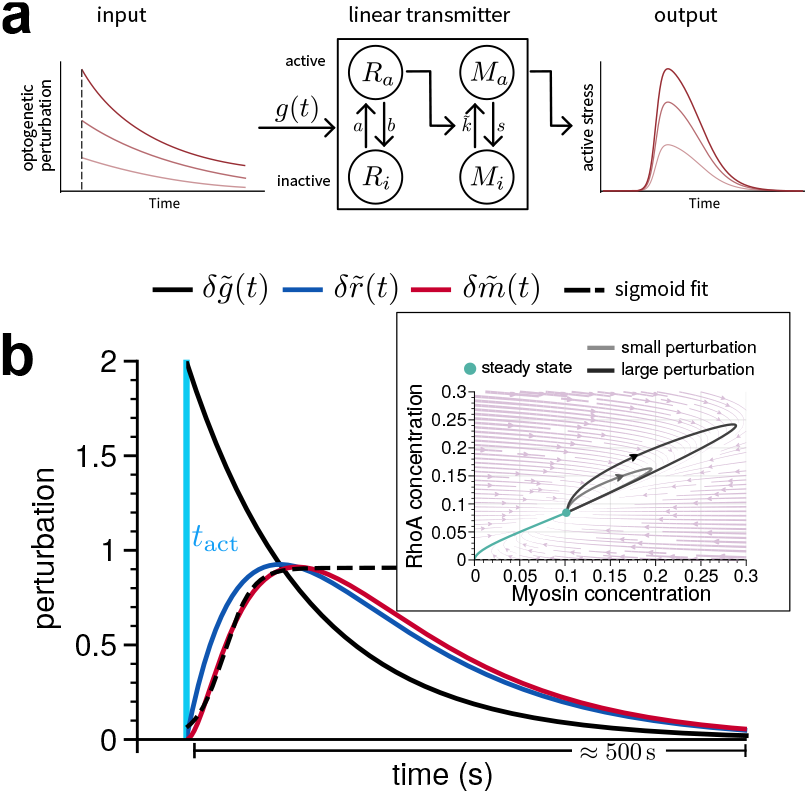
Simple model for the RhoA-pathway. (a) Optogenetic activation of strongly adherent cells often results in a homeostatic response, which is described best by a weakly activated linear signaling cascade. For weak perturbations, the strength of the output signal scales linearly with strength of the input signal. (b) Time course of the normalized concentration perturbations of active RhoA (blue) and myosin (red) after rapid increase of GEF concentration (black) upon photoactivation at *t* = *t*_act_. The dashed line represents a sigmoidal fit to the increasing edge of the myosin concentration in order to estimate the time scale of the increase 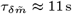. The inset shows the time evolution of the full system of ODEs in the phase-plane. The green line displays the evolution into the only stable fixed point of the system. Two perturbations for *α* = 2 (gray) and *α* = 4 (black) are shown and demonstrate the linearity and scalability of the system.

Here *g*_ss_ represents a normalized steady-state GEF concentration (fraction of active GEF, for *t ≤ t*_act_). After an abrupt light-mediated increase of concentration *g*_a_ at *t* = *t*_act_ (where *H*(*t*) is the Heaviside function), the time course of GEF concentration for *t > t*_act_ follows a decaying exponential.

This input signal consequently triggers a reaction cascade by activating RhoA which in turn activates myosin II. All reactions are modeled by a law of mass action with positive valued activation rate constants *a* and 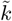. Further we assume that all active components deactivate spontaneously described by the positive valued rate constants *b* and *s* and we express the reaction kinetics as

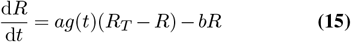

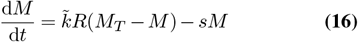

where for brevity we write *R*_*a*_ ≡ *R* and *M*_*a*_ ≡ *M*. Here we additionally assume that the total amount of each signaling component is conserved on the studied time scale, such that concentrations of the inactive species *R*_*i*_ and *M*_*i*_ are given by the difference of the total concentration and the active concentration *R*_*i*_ = *R*_*T*_ *™ R* and *M*_*i*_ = *M*_*T*_ *™ M*. Another simplification is made by considering the limit of a weakly activated signaling cascade (67) for which *R*_*T*_ − *R* ≈ R_T_ and *M*_*T*_ − *M* ≈ M_T_ such that the system can be written as

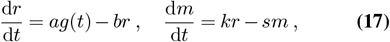

where we divided by the total concentration and hence set *r* = *R/R*_*T*_, *m* = *M/M*_*T*_ and 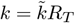. Before photoactivation (*t ≤ t*_act_) we have δ*g*(*t*) = 0 and in this case obtain the stadystate concentrations for RhoA and myosin as

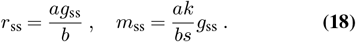

The time evolution after perturbation (*t > t*_act_) may generally be written as

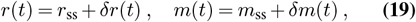

where *δr*(*t*) and *δm*(*t*) denote the time-dependent perturbations of the steady state. Together with Eq. (17) we end up with the time evolution of the perturbation which is given by

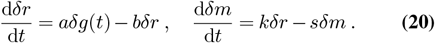

In experiments one usually quantifies the relative activity increase with respect to the activity baseline. We therefore normalize the perturbation with respect to the steady state concentrations and obtain

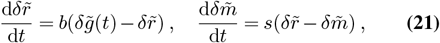

with 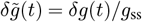 The strength and time course of the relative RhoA and myosin perturbations is controlled by the two deactivation rates *b* and *s* as well as the strength of the input signal *α* ≡ *g*_a_*/g*_ss_ and its decay rate *λ*. For the parametrization of this linearized model we refer the reader to the supplementary text. This system of equations Eq. (21) can be solved analytically for a spatially homogeneous system (solution given in supplemental text). Typical time courses of the perturbations are shown in Fig. 3b, where the inset displays the time evolution of Eq. (17).

## Results

### Optogenetic activation of a cell doublet

We start by simulating the cell doublet, cf. Fig. 2, with linear Rho-signaling as shown in Fig. 3. The corresponding computer code is documented in the supplemental text and the mechanical equations are parametrized according to Table S3. We keep all parameters fixed, except the strength of the strain-dependent feedback 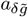 and the viscoelastic time scale *τ*_*c*_. Since we are mainly interested in the response to the perturbation, we omit baseline contractility, i.e. we do not consider a strain in the cell layer before activation. The reference shape was chosen to resemble the typical reference shape of a maturely adherent cell doublet with a vertical oriented cell-cell junction across the symmetry center of the pattern and two pronounced invaginated arcs spanning between the vertical bars of the H-shaped micropattern (52). The cells are assumed to contract isotropically with 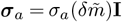.

We optogenetically activate the left cell Ω_*l*_ at time *t* = *t*_act_, compare Fig. 4a. The time evolution of the GEF-perturbation for *t* ≥ t_act_ is given by

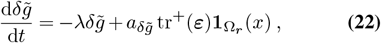

where we include the strain dependent feedback in the right cell via the indicator function **1**_Ω*r*_ (*x*) with **1**_Ω*r*_ (*x*) = 1 if *x* ∈ Ω_*r*_ and **1**_Ω*r*_ (*x*) = 0 otherwise. This means that the strain-dependent feedback is only active in the right cell Ω_*r*_. Having a feedback mechanism in the left cell we can expect non-trivial temporal behavior if positive passive strains build up in the left cell.

**Fig. 4.**
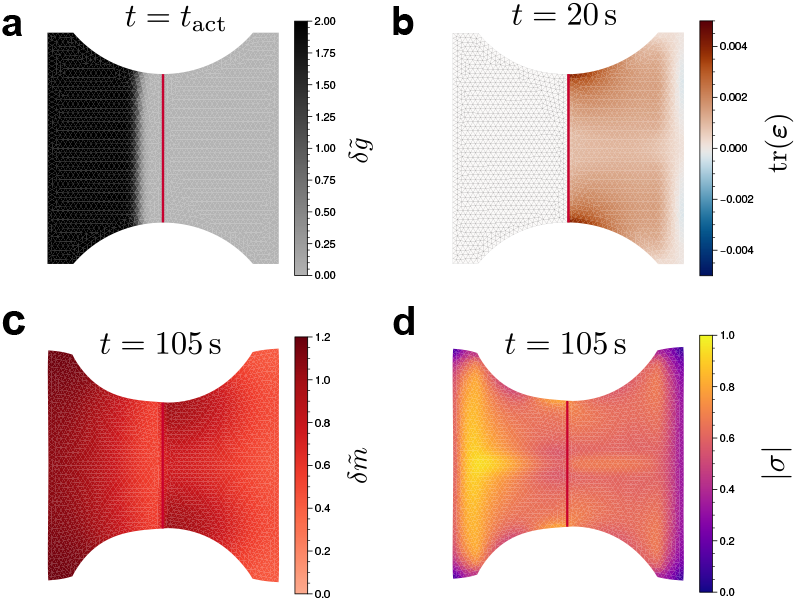
Simulation of the optogenetic activation of a cell doublet in the case of strong coupling. (a) Activated region (in black) and the GEF perturbation 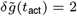. In accordance with experiments, only a fraction of the left cell is illuminated in order to avoid illumination of the right cell. (b) Coupling measure tr ***ε*** in the non-activated cell (right cell) which reaches its maximal value of approx. 5 · 10^−3^ after 20 s. (c) Myosin concentration 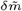 around the time of maximal strain energy. (d) Frobenius norm of the resulting Cauchy stress. The parameters for this simulation can be found in Tables S2 and S3. Further we used 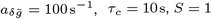 and time step Δt = 0.5 s. Full time sequence shown as Movie S1. Weak coupling shown as Movie S2.

The optogenetic activation at *t*_act_ = 5 s is achieved by setting 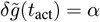. This time point is shown in Fig. 4a with a 100% increase (*α* = 2) of the steady-state GEF concentration upon photoactivation. This GEF perturbation triggers the RhoA-pathway which leads to active contraction in the activated left cell. As contraction in the left cell progresses, passive positive strains are generated in the right cell as it is stretched (Fig. 4b). This stretch leads to activation of GEF and hence triggers a contractile response in the right cell. Fig. 4c and d show the active myosin in both cells as well as the Frobenius norm of the resulting Cauchy stress 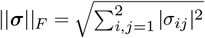, respectively, for a case where parameters are chosen such that both cells deform approximately symmetrically. We note that the active myosin pattern closely follows the pattern of passive strains in the right cell with concentration peaks near the cell periphery at the cellcell junction. Looking at the stress pattern, we further notice that the left cell contracts stronger than the right cell. However, the invaginated arc remains fairly symmetrical. This demonstrates that a visually symmetrical contraction can occur even when both cells do not contract equally strongly, but at different locations within the cell. Here the left cell generates active stresses mainly near the vertical bar of the H-pattern, while the right cell generates active stresses more near the periphery.

To investigate these observations further we define different measures in order to understand the input-output relation between the left and right cell and to quantify the efficiency of the coupling. One way to quantify this is to measure the strain-energy *U*_*s*_ transfered to the substrate

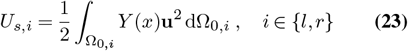

and compare the strain energy deposited on the left and right parts of the pattern (Fig. 5a (left)). Another way is to measure the asymmetry *A* of the contraction by quantifying the shape of the contour *y*(*x*) with respect to the symmetry axis of the pattern and the inward displacement of the cell contour (Fig. 5a (right))

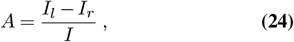

with 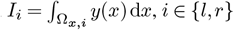.

**Fig. 5.**
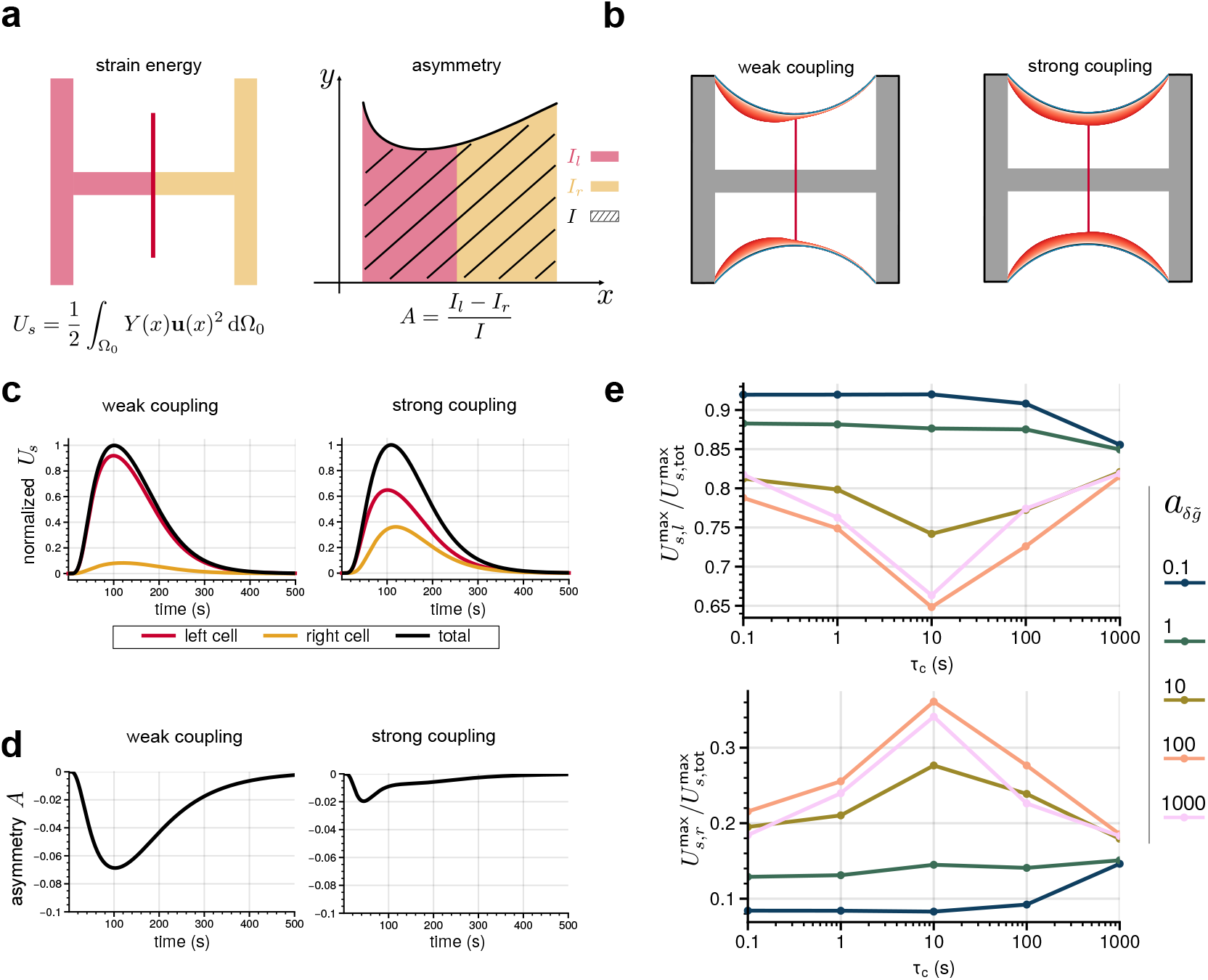
Mechanochemical coupling in cell doublets. (a) One measure for the asymmetry of the process is monitoring the deposited strain energies *U*_*s*_ in the left and right halves of the micropattern, respectively. Another measure is monitoring shape changes through asymmetry *A*, which follows from the integrals under the left and right halves of the contour. (b) Cell shapes in simulations with weak and strong active coupling. Stronger coupling may lead to visually symmetric contraction. Black line represents contour shape at last time step. (c) Effects of weak and strong coupling in terms of strain energy as a function of time. Strain energy curves were normalized with respect to the maximum of the total strain energy curve. (d) Same for shape asymmetry. (e) Efficiency of the coupling defined by the ratio of the relative maxima 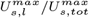 for the left vs. 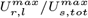 for the right cell, for different combinations of the free parameters, namely coupling strength 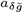 and viscoelastic time scale *τ*_*c*_. For sufficiently large coupling one sees optimal coupling if the viscoelastic time scale corresponds to the time scale of the linear activation cascade.

In Fig. 5b we show the shape of the invaginated arc throughout the whole contraction process. This demonstrates that the symmetry of the contraction is strongly shaped by the coupling described by the rate 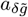. For a stronger coupling (large value of 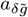) the cell doublet contracts visually symmetrically in comparison to a weaker coupling (small value of 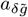), where invagination of the contour is clearly asymmetric and tends towards the activated cell.

In Fig. 5c we show the normalized strain energy *U*_*s*_ as a function of time, where each strain energy is normalized with respect to the maximal total strain energy *U*_*s*,tot_. We observe that the time course of the total strain energy is only slightly changed by the coupling strength 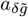. In the case of weak coupling more than 90% of the deposited strain energy is generated by the left cell and the right cell remains almost completely passive. Note that even for 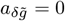 the strain energy on the right hand side of the pattern is expected to be non-zero due to passive deformations by pulling of the left cell. In case of a strong coupling the right cell shows an active response and significantly contributes to the overall generated strain energy. These observations are confirmed by a quantification of the contour deformation as a function of time. For the weak coupling the contour remains strongly asymmetric throughout the majority of the simulation time. For the strong coupling asymmetry is only present for small times until the right cell actively pulls back and symmetrizes the periphery (Fig. 5d).

Until now we only altered the rate 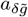 and kept the viscoelastic time scale *τ*_*c*_ constant. We next addressed the question of how the viscoelastic properties of the cell influence the efficiency of the coupling and therefore simulated several combinations of 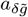 and *τ*_*c*_ by varying both parameters over several orders of magnitudes. We consider the coupling to be efficient, if the ratio 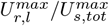 is high. The results are displayed in Fig. 5e.

We find that coupling efficiency is positively correlated with the coupling strength 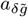, provided the viscoelastic time scale is such that stresses do relax on a comparable time scale. As a function of the viscoelastic time scale we notice that coupling efficiency increases with an increasing viscoelastic time scale, reaches an optimum around intermediate values and decreases again for larger values of of *τ*_*c*_. The optimal value for *τ*_*c*_ is found to be at *τ*_*c*_ = 10 s, which is in the same order of magnitude as experimentally measured values (50, 68). A sigmoidal fit to the time course of the myosin concentration (Fig. 3b) yields a time scale of 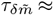 11 s similar to the viscoelastic time scale. This result suggests that for an efficient mechanochemical coupling the viscoelastic time scale, i.e. the time scale at which the cell can react to mechanical stimuli, has to be in tune with the time scale at which the full linear cascade can react to those. In case of very large *τ*_*c*_ the left cell cannot deform sufficiently to trigger a response in the right cell. For small values of *τ*_*c*_ the right cell starts to counteract deformations quickly and does not remain in a stretched state for a sufficiently long time to allow the coupling mechanism to unfold its action.

### Optogenetically stimulated contraction wave in a chain of cells

Having established a framework for the mechanochemical interplay of two cells, we can generalize to more cells, for example a chain of cells on a micropatterned line (compare Fig. 1b). It is the strength of our discontinuous Galerkin approach that now such simulations are easy to implement and very efficient. The cell chain is realized computationally by connecting equally sized cells along the x-axis. Each cell now adheres in a homogeneous fashion to an elastic substrate. In contrast to the doublet, we here include a mechanochemical coupling also for the activated cell, thus we allow for the possibility that a wave is reflected at the right and comes back to the left. Like for the cell doublet, we again assume that due to strong cell-matrix adhesion, the Rho-system has a stable fixed point and that the simple model is sufficient. To model that the cells tend to polarize when placed on lines, we use a unidirectional active stress tensor such that cells only contract in longitudinal direction of the line of cells 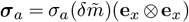. All parameters used in the simulation are given in Tables S2 and S4.

We now activate the left cell at *t* = *t*_act_. The time evolution of the GEF-concentration for *t > t*_act_ is then described by

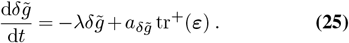

As for the previous study of the cell doublet, we again vary the parameters *τ*_*c*_ and 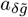 to investigate the response of the system. Depending on parameters, we now observe three different responses: (1) non-transmissive (the stimulus dies out); (2) transmissive wave propagation, i.e. the most right cell is activated once; and (3) oscillatory waves going persistently through the system.

Fig. 6a depicts the displacement field for the different states.

**Fig. 6.**
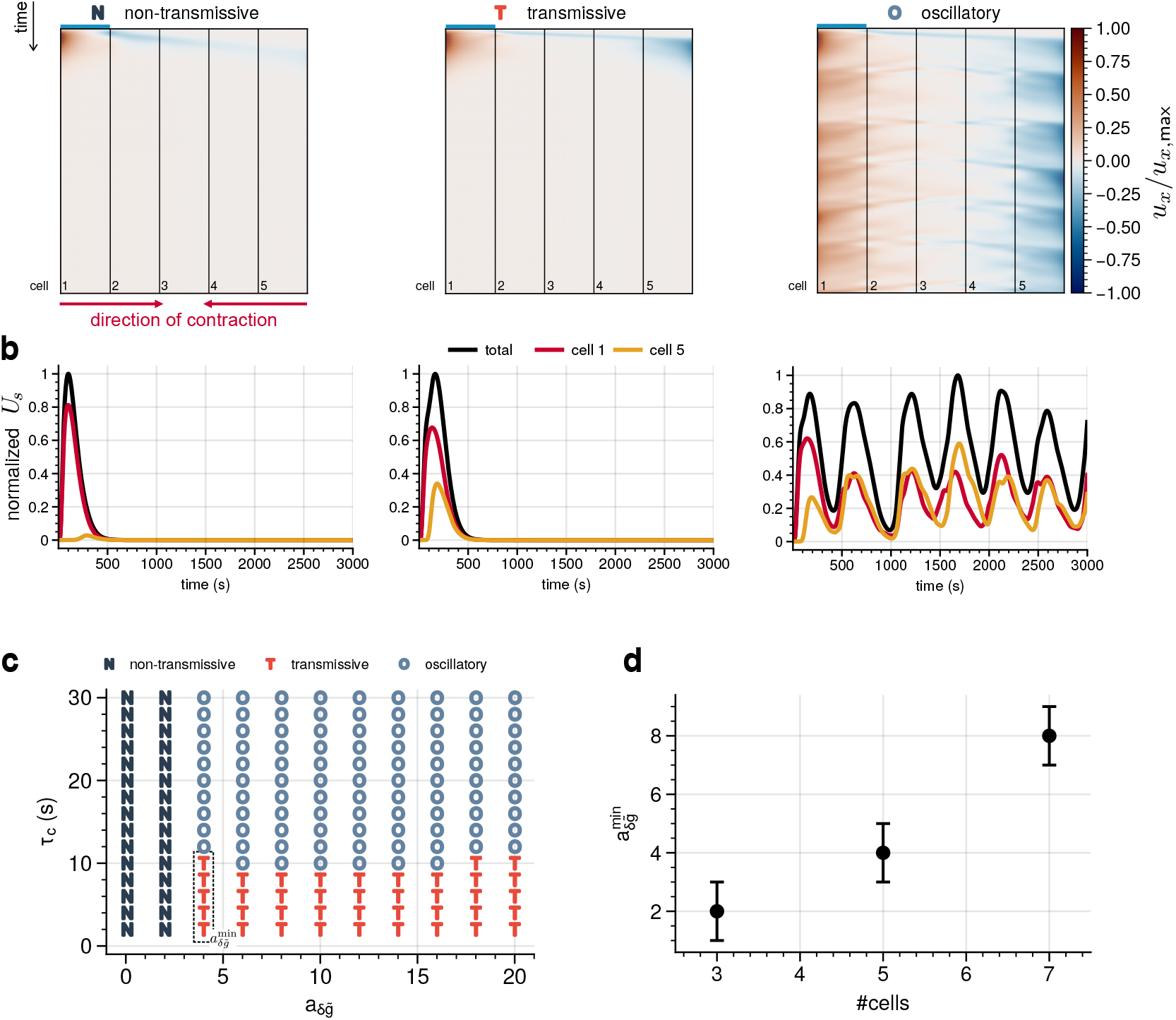
Mechanochemical coupling in a chain of cells. (a) Kymographs of the substrate displacement field normalized to the maximal displacement for five cells in a row and for three different conditions. At *t* = 5 s cell 1 (left) is activated (blue bar). The contraction of the activated cell initiates a contraction wave propagating from cell 1 (left) to cell 5 (right). Depending on the parameters we observe non-transmissive (N), transmissive (T) or oscillatory states (O). For sufficiently large strain-dependent feedback 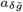 and sufficiently large viscoelastic time scale *τ*_*c*_, self-sustained oscillations emerge. (b) Time evolution of the strain energy for the activated cell 1 and cell 5, i.e. the last cell of the line, corresponding to the kymographs given in (a). (c) Phase diagram as a function of activation strength and viscoelastic times scale. Strong coupling and large viscoelastic times are required for oscillations. (d) Effect of cell number. The transition from non-transmissive to transmissive occurs at larger values of 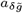 with increasing length of the cell chain (error bars highlight discrete parameter sampling). A full simulation is given as Movie S3.

Note that the displacement field is directly correlated with the generated traction forces through **T** = *Y* **u**. In the case of a non-transmissive parameter regime, i.e. for small 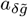, the contraction signal strongly decays during propagation from cell 1 (left) to cell 5 (right) and no substantial traction forces are generated at the right end of the line of cells, see Fig. 6a on the left. This observation has been quantified by comparing the substrate strain energies generated by cell 1 and 5 as a function of time, see Fig. 6b on the left.

With increasing values of 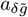, the transmission of the signal becomes more and more efficient and the active response of cell 5 comparable to cell 1, see the middle panels of Fig. 6a and b. Since there is no absolute definition of effective transmission, a threshold criterion must be chosen and we define the system as transmissive if 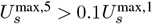.

Finally, varying also the viscoelastic time scale of the system, we observe persistent oscillatory states in the parameter regime of sufficiently large 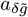 and *τ*_*c*_, see the right panels of Fig. 6a and b. The frequency of the oscillations depends strongly on the combination of 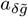 and *τ*_*c*_ and can be further classified into burst-like oscillations and contractile oscillations. The former correspond to large variations in strain energy, while the latter relate to small fluctuations around an elevated increased strain energy plateau (see Fig. S2).

Fig. 6c shows the phase diagram as a function of the coupling strength and the viscoelastic time scale. For a strongly coupled system (i.e. sufficiently large 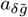), the transition between transmissive and oscillatory states occurs around *τ*_*c*_ ≈ 10 s, which again coincides with the time scale of the myosin relaxation. The number of cells in the line is also relevant for the signal transmission. Fig. 6d shows that the transition from non-transmissive to transmissive mechanochemical signaling needs larger values of 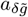 when increasing the length of the cell chain. This emphasizes that mechanochemical signaling tends to be of dissipative nature, and hence a cell can reach out only to a limited number of other cells, except if a strong amplification exists, like in action potentials.

### Cell monolayer with non-linear Rho-pathway

We finally turn to a cell monolayer (compare Fig. 1b). Again the numerical implementation is relatively easy given our discontinuous Galerkin approach and we demonstrate this here for 28 hexagonally arranged cells. In contrast to the doublet and the line of cells, cell-matrix adhesion is now less relevant and we do not assume that the cells in the layer are polarized in the lateral direction. Moreover in the two-dimensional case, cells have many neighbors and therefore many inputs. These aspects suggest that the simple linear chain model for the Rho-pathway might not be sufficient anymore and that more complex dynamics might arise, similar to the excitable dynamics known from the cortex of oocytes (31, 33). We therefore now use the full non-linear model for the Rho-pathway as established earlier by Kamps et al. (34). The corresponding kinetic equations are given in Eqs. (S29-S31) (supplemental text). As investigated in Kamps et al. (34), they exhibit Turing-type instabilities resulting from fast (inactive) and slow (active) diffusing species paired with positive and negative feedback loops of activator-inhibitor type. Here we have chosen the parameters for the RD-system such that traveling waves can form.

The cell monolayer is shown in Fig. 7a with intercellular junctions shown in red and the color code depicting the levels of active stress/active myosin. The cell in the middle on the left hand side of the monolayer was activated by inducing an instability through small random fluctuations in the GEF- and RhoA-concentrations (the same effect can be obtained by optogenetic activation, but for the non-linear model, the exact mode of activation is less relevant). Several distinct contraction peaks form and the induced deformations slowly activate the RhoA-pathway and allow the active stresses to spread through the tissue, as indicated by the white arrows in Fig. 7a. The initially weak and rather uniform contractions eventually turn into strong and more localized traveling contraction waves, see Fig. 7b, until the whole monolayer is strongly activated, see Fig. 7c. For longer times, a dynamic steady state is reached, where the deformations are most prominent near the free edges, see Fig. 7d.

**Fig. 7.**
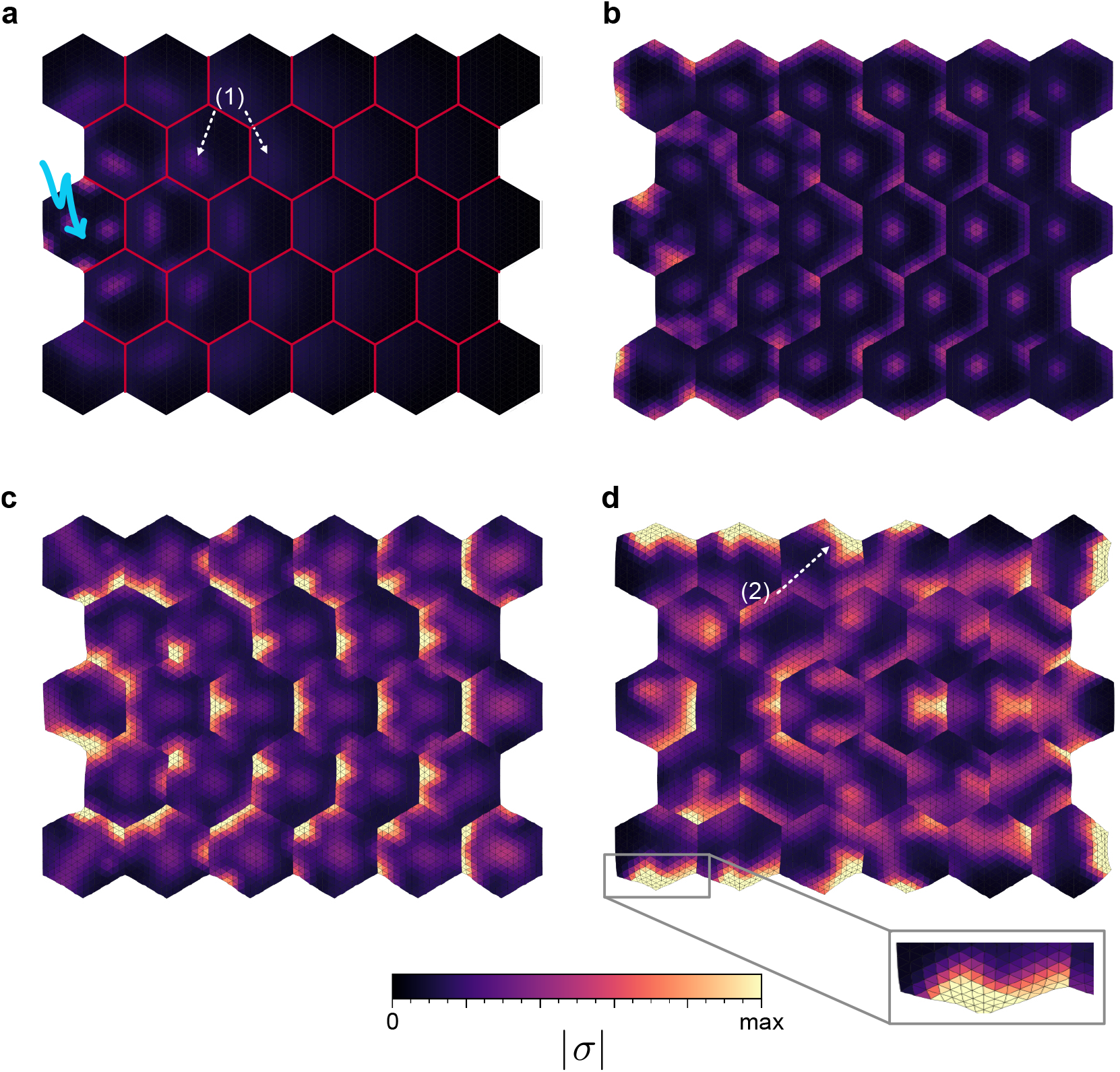
Mechanochemical coupling in a cell monolayer with a non-linear model for the Rho-pathway. (a) Total stress in the tissue shortly after inducing a spatio-temporal contraction pattern in the centered cell on the left side of the tissue. This initial perturbation spreads by means of the mechanochemical feedback through the tissue (white arrows). (b) and (c) Tissue at a later stage, where the initially small perturbations have developed into substantial contraction waves in each of the cells. (d) After transient dynamics, prominent deformations are visible in the free edges of the cell layer (white arrow and inset). Parameters can be found in Table S1 and Table S5. Full dynamics shown as Movie S4.

## Discussion and Conclusion

Here we have shown that the discontinuous Galerkin (DG) finite element method (FEM) is ideally suited to model mechanochemical coupling in cell layers. Because both the reaction-diffusion (RD) and the continuum mechanical models lead to partial differential equations (PDEs), finite elements are the natural approach to couple them in a numerically efficient framework. The DG method in addition is ideal to describe the effect of concentration jumps at the cell-cell boundaries while still keeping the continuous nature of the PDEs for the RD-system. In contrast, the PDEs for the mechanics can be treated well with standard continuous Galerkin (CG), because displacements are continuous across the boundaries. Our approach is very general both in terms of biochemistry (as exemplified by the linear regulation cascade versus the fully nonlinear RD-system) and the material law (here a linear Kelvin-Voigt-type material). Motivated by the homeostatic nature of adherent cell layers, it is however centered on elastic systems. For flowing systems, which are typically modeled with active gel theory (24–28), one had to switch from the Lagrangian to the Eulerian framework.

To couple the RD- and mechanical sub-systems, we used very simple but reasonable assumptions. Like in active gel theory, active stress is coupled to the concentration of myosin II (the last molecule of the signaling cascade), and the activation rate at the beginning of the signaling cascade (typically an exchange factor that can be controlled by optogenetics) is assumed to be proportional to the positive part of the trace of the strain tensor. In future work, one could also consider non-linear model versions both for the material law (e.g. Neo-Hookean) and the coupling (e.g. higher order invariants of the strain tensor).

We benchmarked our approach by investigating in detail the optogenetic activation of a cell doublet on a H-shaped micropattern, as recently studied experimentally in Ref. (52). Because that work did not include any experiments on signal transduction, we cannot compare our results directly to this work. However, our work shows that the detailed RD-model used here can explain the experimental observation that mechanical interactions between an optogenetically activated cell and a neighboring non-activated cell can trigger an active contractile response. Interestingly, our simulations further predict that a cell doublet can exhibit an apparently symmetric contraction even when the left and right cells display distinct internal active stress distributions. This also shows that shape is not a unique readout of the internal state of the cell and that care has to be applied when estimating cell forces from shape.

We then exemplified the scalability of our DG-approach to larger systems by looking at force and signal transmission via contraction waves in chains of cells, similar to recent experiments on optogenetic activation of cell trains (53). In these two cases, cell-matrix adhesion is very strong and cells tend to be homeostatic, which means that they return to baseline after optogenetic stimulation (50). Therefore we again used the simple model of a linear chain for the Rho-signaling pathway. Again a direct comparison to the experimental data is not possible because the signaling part was not investigated, but on a qualitative level, we find similar results, in particular traction force patterns with local reversals in force direction near the center of the cell train and that larger trains attenuate signal propagation.

We finally addressed the case of mechanochemically excitable monolayers, which are known to develop more complex dynamic patterns, possibly because cell-cell adhesion dominates and cells are less polarized. In particular, experiments have found that wave propagation is rather common in such systems (7, 9, 10) and recent simulation have shown that this should also apply to three dimensions (11). In this case, we therefore used a more comprehensive and non-linear model for the Rho-pathway (34). We then observed excitable dynamics that does not die down again, similar to experimental observations in the cortex of oocytes (31, 33), but also adherent cells with intermediate activation of the Rho-pathway (34).

In all of these cases, we identified conditions under which mechanochemical signaling leads to strong propagation of the signal from one cell to the other, and even to wave propagation. We find that force transmission is best when the viscoelastic time scale of the cell and the time scale of relaxation of the Rho-pathway after activation are of the same order. This finding makes sense because for the bidirectional feedback to work, both processes need to work on similar time scales. The numerical value of 10 s also makes sense because this is the time scale on which adhesion contacts can remodel. We note that when elastic effects have to be faster, e.g. in the venus trap or in the mechanochemical cycle of molecular motors, the system tends to separate the slow buildup of elastic energy and its fast release. Here, however, such a separation is not needed and in fact would be a disadvantage. As shown here, the mechanochemical coupling between neighboring cells can lead to long-ranged and even wave-like signal propagation, as e.g. recently measured by optogenetic activation of the developing zebrafish neural rod (69). This agrees with earlier work on optogenetic activation in Drosophila, which showed that local activation can affect global processes like gastrulation (70, 71).

For the cell doublet, we used the coupling to an elastic foundation as a readout of the mechanical coupling of the two neighboring cells, in addition to the asymmetry in shape. In the future, the coupling to the elastic foundation could be replaced by a continuum substrate; then one could also model the effect of mechanical cell-cell communication through the substrate, which would be a natural extension of our approach. On the biochemical side, it would be interesting to go beyond modelling of the Rho-pathway and also include the effect of other known signaling molecules, including Rac/Cdc42, ERK and merlin. The numerical DG FEM framework established here now opens the door to quickly explore the effect of such pathways, and thus to establish a multiscale modeling framework that connects molecular processes to effective systems behavior.

## Data and code availability

The Python code for the examples described above is publicly available at https://github.com/dworthmuller/MechanochemicalCoupling.

## Acknowledgments

U.S.S. wishes to express his gratitude to Erich Sackmann for many inspiring discussions on the physics of cells. D.W. and U.S.S. acknowledge funding through the DFG (Deutsche Forschungsgemeinschaft) grant MechanoSwitch SCHW 834/2-1. F.Z. and U.S.S. were supported by the cluster of excellence STRUCTURES funded by the DFG (EXC 2181/1 – 390900948). D.W. received funding from a European Research Council (ERC) grant ERC-SyG 101071793, awarded to Pierre Sens.

## Author Contributions

All authors designed the research together. D.W. performed the research. F.Z. and U.S.S. supervised the project. D.W. and U.S.S. wrote the original draft of the paper. All authors reviewed and approved the paper.

## Supplementary Material

An online supplement to this article can be found by visiting BJ Online at http://www.biophysj.org.

