## Supplemental file for "Modelling mechanochemical coupling in optogenetically activated cell layers"

### - Supplemental text -

D. Wörthmüller, F. Ziebert and U.S. Schwarz

#### General notation

In solid mechanics one typically expresses model equations in the Lagrangian (material) framework by describing a deforming solid in terms of coordinates $\hat{\mathbf{x}}$ representing the position of material particles in the undeformed configuration $\Omega_0$. The current position of the material particles in the deformed configuration $\Omega(t)$ at time $t$ are given by $\mathbf{x} = \boldsymbol{\chi}(\hat{\mathbf{x}}, t)$. The two configurations are connected by the displacement vector field $\hat{\mathbf{u}}(\hat{\mathbf{x}}, t) = \mathbf{x}(\hat{\mathbf{x}}, t) - \hat{\mathbf{x}}$. The deformation gradient tensor is defined as $\hat{\mathbf{F}} = \partial \mathbf{x}/\partial \hat{\mathbf{x}} = \mathbf{I} + \hat{\nabla}\hat{\mathbf{u}}$ and measures the local change of relative position of two points at the transition from the undeformed to the deformed configuration. Its determinant $\hat{J} = \det(\hat{\mathbf{F}})$ represents the local volume change. Describing the governing equations in terms of the coordinates $\mathbf{x}$ is known as the Eulerian framework where a fixed point in space is observed. In the limit of linear elasticity we do not distinguish between Lagrangian and Eulerian description and drop the $\wedge$-symbol.

#### Reaction-diffusion on time dependent domains in Lagrangian frame of reference

For the pull-back of

$$\frac{\partial c_i}{\partial t} + \nabla \cdot (\mathbf{v}(\mathbf{x}, t) c_i) = \nabla \cdot (\mathbf{D} \nabla c_i)) + R_{c_i}(t) \tag{S1}$$

to the reference configuration (Lagrangian coordinates) we exploit the transformation rules for scalar and vector fields as well as the involved differential operators (1). For mappings between the two reference systems we use the motion function $\mathbf{x} = \boldsymbol{\chi}(\hat{\mathbf{x}}, t)$. Each Eulerian field has a Lagrangian counterpart which states the equivalence of the two descriptions. Hence we write $c(\mathbf{x}, t) = \hat{c}(\hat{\mathbf{x}}, t)$ for the concentration field and $\mathbf{v}(\mathbf{x}, t) = \hat{\mathbf{v}}(\hat{\mathbf{x}}, t)$ for the velocity field. Here $\hat{\mathbf{v}} = \partial_t \boldsymbol{\chi}(\hat{x}, t)$ denotes the material velocity. For the gradient of the scalar field $c(\mathbf{x}, t)$ it holds

$$\nabla c = \hat{\mathbf{F}}^{-\mathsf{T}} \hat{\nabla} \hat{c}\,. \tag{S2}$$

For the gradient of the vector field $\mathbf{v}$ we have

$$\nabla \mathbf{v} = \hat{\nabla} \hat{\mathbf{v}} \hat{\mathbf{F}}^{-1} \; . \tag{S3}$$

The divergence of the vector field transforms according to

$$\nabla \cdot \mathbf{v} = \frac{1}{\hat{J}} \hat{\nabla} \cdot (\hat{J} \hat{\mathbf{F}}^{-1} \hat{\mathbf{v}}) \; , \tag{S4}$$

and we note that Piola's identity is given by

$$\hat{\nabla} \cdot (\hat{J} \hat{\mathbf{F}}^{-1}) = 0 \; . \tag{S5}$$

The material time derivative of a field expressed in Eulerian coordinates is given by the convective derivative

$$\frac{\mathrm{d}}{\mathrm{d}t} c(\mathbf{x}, t) = \frac{\partial}{\partial t} c + \mathbf{v} \cdot \nabla c \equiv \frac{\partial \hat{c}}{\partial t} \tag{S6}$$

Starting from the reaction-diffusion equation in Eulerian coordinates

$$\frac{\partial c_i}{\partial t} + \mathbf{v} \cdot \nabla c_i + c_i (\nabla \cdot \mathbf{v}) = \nabla \cdot (\mathbf{D} \nabla c_i)) + R_{c_i}(t) \; , \tag{S7}$$

we note that the first two terms correspond to the convective time derivative. The third term on the left and the first term on the right hand side of the equation can be replaced by using the Piola transform Eq. (S4) such that

$$\frac{\partial \hat{c}_i}{\partial t} + \hat{c}_i \frac{1}{\hat{J}} \hat{\nabla} \cdot (\hat{J} \hat{\mathbf{F}}^{-1} \hat{\mathbf{v}}) = \frac{1}{\hat{J}} \hat{\nabla} \cdot (\hat{J} \hat{\mathbf{F}}^{-1} \mathbf{D} \hat{\mathbf{F}}^{-\mathsf{T}} \hat{\nabla} \hat{c}_i)) + R_{\hat{c}_i}(t) \; , \tag{S8}$$

and after multiplying by $\hat{J}$ we obtain

$$\hat{J} \frac{\partial \hat{c}_i}{\partial t} + \hat{c}_i \hat{\nabla} \cdot (\hat{J} \hat{\mathbf{F}}^{-1} \hat{\mathbf{v}}) = \hat{\nabla} \cdot (\hat{J} \hat{\mathbf{F}}^{-1} \mathbf{D} \hat{\mathbf{F}}^{-\mathsf{T}} \hat{\nabla} \hat{c}_i)) + \hat{J} R_{\hat{c}_i}(t) \; . \tag{S9}$$

Applying Piola's identity to the second term on the left we find

$$\hat{c}_i \hat{\nabla} \cdot (\hat{J} \hat{\mathbf{F}}^{-1} \hat{\mathbf{v}}) = \hat{c}_i \hat{J} \hat{\mathbf{F}}^{-1} \hat{\nabla} \hat{\mathbf{v}} = \hat{c}_i \hat{J} \hat{\mathbf{F}}^{-1} \frac{\partial \hat{\mathbf{F}}}{\partial t} = \hat{c}_i \frac{\partial \det(\hat{\mathbf{F}})}{\partial t} = \hat{c}_i \frac{\partial \hat{J}}{\partial t} \; , \tag{S10}$$

and further using $\hat{\mathbf{C}} = \hat{\mathbf{F}}^{\mathsf{T}} \hat{\mathbf{F}}$ we end up with

$$\frac{\partial}{\partial t} (\hat{J} \hat{c}_i) = \hat{\nabla} \cdot (\hat{J} \mathbf{D} \hat{\mathbf{C}}^{-1} \hat{\nabla} \hat{c}_i) + \hat{J} R_{\hat{c}_i}(t) \; . \tag{S11}$$

### Finite-Element formulation

#### Symmetric Weighted Interior Penalty Discontinuous Galerkin method

The interface condition on the cell-cell junction

$$-\hat{J} \mathbf{D} \hat{\mathbf{C}}^{-1} \hat{\nabla} \hat{c}_i \cdot \mathbf{N}_{\mathsf{CCJ}} = 0 \quad \text{on} \quad \Gamma_{\mathsf{CCJ}} \; , \tag{S12}$$

leads to a discontinuity in the solution. In order to account for abrupt concentration changes from one subdomain to the other we need a method which allows discontinuous functions across the membrane. The standard choice for such a problem is the discontinuous Galerkin method. In contrast to the continuous Galerkin methods, continuity and smoothness of the involved DG-functions is only enforced element-wise such that the solution may be discontinuous across element boundaries. If and where desired, continuity can be enforced through an appropriate penalty term. These methods are known as interior penalty discontinuous Galerkin methods (IPDG) (2–4). However, as can be seen from Eq. (S11) the diffusivity $\boldsymbol{\alpha} \equiv \hat{J}\mathbf{D}\hat{\mathbf{C}}^{-1}$ in the Lagrangian frame is position dependent due to local deformations of the domain. Hence, we are dealing with heterogeneous diffusion and use a Symmetric Weighted Interior Penalty (SWIP) discontinuous Galerkin scheme (5). Let $\mathcal{T}(\Omega_0)$ be the triangulation of the domain $\Omega$ into finite elements $e \in \mathcal{T}(\Omega_0)$. Further, let $\mathcal{F}$ denote the union of the boundary facets of all elements $e$. We distinguish between external facets $\mathcal{F}_{\text{ext}}$, internal facets $\mathcal{F}_{\text{int}}$ and membrane facets $\mathcal{F}_{\text{M}}$ such that $\mathcal{F} = \mathcal{F}_{\text{ext}} \cup \mathcal{F}_{\text{int}} \cup \mathcal{F}_{\text{M}}$ with $\mathcal{F}_{\text{int}} = \mathcal{F} \setminus (\mathcal{F}_{\text{ext}} \cup \mathcal{F}_{\text{M}})$. Next, by $\hat{c}_-$ and $\hat{c}_+$ we denote scalar valued functions on two neighboring elements $e_-$ and $e_+$. The normal vectors on a common facet of $e_\pm$ are given by $\mathbf{N}_\pm$. For example, $\mathbf{N}_-$ defines the outward directed normal on $e_-$ pointing into $e_+$. Following the SWIP-DG notations we introduce the jump and the weighted average of a quantity as $[\![\hat{c}]\!] \equiv \hat{c}_+\mathbf{N}_+ + \hat{c}_-\mathbf{N}_-$ and $\{\hat{c}\}_\omega \equiv \omega_+\hat{c}_+ + \omega_-\hat{c}_-$, respectively. Analogously, for piecewise vector valued functions $\hat{\mathbf{q}}$ one defines jump and weighted average as $[\![\hat{\mathbf{q}}]\!] \equiv \hat{\mathbf{q}}_+\mathbf{N}_+ + \hat{\mathbf{q}}_-\mathbf{N}_-$ and $\{\hat{\mathbf{q}}\}_\omega \equiv \omega_+\hat{\mathbf{q}}_+ + \omega_-\hat{\mathbf{q}}_-$, respectively. The weights are defined as $\omega_\pm = \delta_\alpha^\mp/(\delta_\alpha^+ + \delta_\alpha^-)$, where $\delta_\alpha^\mp$ is obtained from the diffusivity on two neighbouring elements $e_\mp$ by calculating $\delta_\alpha^\mp = \tilde{\mathbf{N}}_e^\mathsf{T}\boldsymbol{\alpha}_\mp\tilde{\mathbf{N}}_e$. Note that the weights fulfill $\omega_+ + \omega_- = 1$. Since we are only interested in having a discontinuity across the membrane facets $\mathcal{F}_{\text{M}}$ we introduce a diffusion dependent penalty term to penalize jumps across all other internal facets $\mathcal{F}_{\text{int}}$ which is defined as the harmonic mean $\gamma_\alpha = 2\delta_\alpha^+\delta_\alpha^-/(\delta_\alpha^+ + \delta_\alpha^-)$. Moreover, one may use these definitions to prove the identity

$$[\![\hat{\mathbf{q}}\hat{c}]\!] = [\![\hat{\mathbf{q}}]\!]\{\hat{c}\}_\omega + \{\hat{\mathbf{q}}\}_\omega[\![\hat{c}]\!] \; . \tag{S13}$$

In the first step of the derivation of the DG weak form we multiply Eq. (S11) with a suitable test function $v_c \in \mathcal{V}$ and integrate over the whole simulation domain $\Omega_0$ which gives

$$\begin{aligned} \int_{\Omega_0} \frac{\partial}{\partial t}(\hat{J}\hat{c})v_c \; \mathrm{d}\Omega_0 - \int_{\Omega_0} \hat{\nabla}\cdot\left(\boldsymbol{\alpha}\hat{\nabla}\hat{c}\right) v_c \; \mathrm{d}\Omega_0 \\ - \int_{\Omega_0} \hat{J}R_c(t)v_c \; \mathrm{d}\Omega_0 = 0 \; . \end{aligned} \tag{S14}$$

Instead of directly using partial integration on the middle term of Eq. (S14) we first split it into a sum over element integrals and then apply Green's first theorem to obtain

$$\begin{aligned} \int_{\Omega_0} \hat{\nabla}\cdot\left(\boldsymbol{\alpha}\hat{\nabla}\hat{c}\right) v_c \; \mathrm{d}\Omega_0 &= \sum_{e\in\mathcal{T}(\Omega_0)} \int_e \hat{\nabla}\cdot\left(\boldsymbol{\alpha}\hat{\nabla}\hat{c}\right) v_c \; \mathrm{d}\Omega_0 \\ &= \sum_{f_e\in\mathcal{F}(\Omega_0)} \int_{f_e} \boldsymbol{\alpha}\hat{\nabla}\hat{c}\cdot\tilde{\mathbf{N}}_e v_c \; \mathrm{d}s \\ &\quad - \sum_{e\in\mathcal{T}(\Omega_0)} \int_e \boldsymbol{\alpha}\hat{\nabla}\hat{c}\cdot\hat{\nabla}v_c \; \mathrm{d}\Omega_0 \; . \end{aligned} \tag{S15}$$

Here, $f_e$ denotes the facets of element $e$ and $\tilde{\mathbf{N}}_e$ describes the outward directed normal vector on the facets of the element. The first term in Eq. (S15) is split again into the exterior, interior and membrane facets

$$\sum_{f_e \in \mathcal{F}(\Omega_0)} \int_{f_e} \boldsymbol{\alpha} \hat{\nabla} \hat{c} \cdot \tilde{\mathbf{N}}_e v_c \; \mathrm{d}s = \sum_{f_e \in \mathcal{F}_{\mathsf{ext}}(\Omega_0)} \int_{f_e} \boldsymbol{\alpha} \hat{\nabla} \hat{c} \cdot \tilde{\mathbf{N}}_e v_c \; \mathrm{d}s + \sum_{f_e \in \mathcal{F}_{\mathsf{int}}(\Omega_0)} \int_{f_e} \boldsymbol{\alpha} \hat{\nabla} \hat{c} \cdot \tilde{\mathbf{N}}_e v_c \; \mathrm{d}s + \sum_{f_e \in \mathcal{F}_{\mathsf{M}}(\Omega_0)} \int_{f_e} \boldsymbol{\alpha} \hat{\nabla} \hat{c} \cdot \tilde{\mathbf{N}}_e v_c \; \mathrm{d}s \; . \tag{S16}$$

Note that each internal facet and each membrane facet is shared by two adjacent elements $e_-$ and $e_+$ (see Fig. S1) such that integrals along the common facets add up to a jump

$$\int_{f_\pm} \boldsymbol{\alpha} \hat{\nabla} \hat{c} \cdot \tilde{\mathbf{N}}_\pm v_c \; \mathrm{d}s = \int_f (\delta_\alpha^+ \hat{\nabla} \hat{c}_+ v_{c,+} - \delta_\alpha^- \hat{\nabla} \hat{c}_- v_{c,-}) \cdot \tilde{\mathbf{N}}_+ \; \mathrm{d}s = \int_f [\![ \boldsymbol{\alpha} \hat{\nabla} \hat{c} v_c ]\!] \; \mathrm{d}s \; . \tag{S17}$$

Summing up over all elements $e$ in Eq. (S15) and Eq. (S16) while respecting zero-flux boundary conditions yields

$$\int_{\Omega_0} \hat{\nabla} \cdot \left( \boldsymbol{\alpha} \hat{\nabla} \hat{c} \right) v_c \; \mathrm{d}\Omega_0 = - \int_{\Omega_0} \boldsymbol{\alpha} \hat{\nabla} \hat{c} \cdot \hat{\nabla} v_c \; \mathrm{d}\Omega_0 + \int_{\mathcal{F}_{\mathsf{int}}} [\![ \boldsymbol{\alpha} \hat{\nabla} \hat{c} v_c ]\!] \; \mathrm{d}s \; . \tag{S18}$$

The last term in Eq. (S18) is further expanded using the identity in Eq. (S13) which yields

$$\int_{\mathcal{F}_{\mathsf{int}}} [\![ \boldsymbol{\alpha} \hat{\nabla} \hat{c} v_c ]\!] \; \mathrm{d}s = \int_{\mathcal{F}_{\mathsf{int}}} [\![ \boldsymbol{\alpha} \hat{\nabla} \hat{c} ]\!] \cdot \{ v_c \}_\omega \; \mathrm{d}s + \int_{\mathcal{F}_{\mathsf{int}}} \{ \boldsymbol{\alpha} \hat{\nabla} \hat{c} \}_\omega \cdot [\![ v_c ]\!] \; \mathrm{d}s \; . \tag{S19}$$

Since the exact solution of the diffusion equation is expected to be smooth we enforce continuity of the fluxes by setting $[\![ \boldsymbol{\alpha} \hat{\nabla} \hat{c} ]\!] = 0$. To further enforce continuity of the solution we exploit $[\![ \hat{c} ]\!] = 0$ and add a term to symmetrize the problem. Additionally, we ensure stability of the problem by adding a stabilizing term according to Ern et al. (5) and Douglas and Dupont (6) which finally leads to

$$\int_{\mathcal{F}_{\mathsf{int}}} [\![ \boldsymbol{\alpha} \hat{\nabla} \hat{c} v_c ]\!] \; \mathrm{d}s = \int_{\mathcal{F}_{\mathsf{int}}} \{ \boldsymbol{\alpha} \hat{\nabla} \hat{c} \}_\omega \cdot [\![ v_c ]\!] \; \mathrm{d}s + \int_{\mathcal{F}_{\mathsf{int}}} \{ \boldsymbol{\alpha} \hat{\nabla} v_c \}_\omega \cdot [\![ \hat{c} ]\!] \; \mathrm{d}s - \int_{\mathcal{F}_{\mathsf{int}}} \frac{s_N}{h} \gamma_\alpha [\![ \hat{c} ]\!] \cdot [\![ v_c ]\!] \; \mathrm{d}s \; . \tag{S20}$$

In Eq. (S20), $s_N$ denotes the so-called Nitsche paramater, which must be chosen sufficiently large to ensure continuity across internal facets (7), and $h$ the average element diameter. Next, we define

$$\mathcal{D}(\hat{c}, v_c, \boldsymbol{\alpha}) := \int_{\Omega_0} \boldsymbol{\alpha} \hat{\nabla} \hat{c} \cdot \hat{\nabla} v_c \; \mathrm{d}\Omega_0 - \int_{\mathcal{F}_{\mathsf{int}}} \{ \boldsymbol{\alpha} \hat{\nabla} \hat{c} \}_\omega \cdot [\![ v_c ]\!] \; \mathrm{d}s - \int_{\mathcal{F}_{\mathsf{int}}} \{ \boldsymbol{\alpha} \hat{\nabla} v_c \}_\omega \cdot [\![ \hat{c} ]\!] \; \mathrm{d}s + \int_{\mathcal{F}_{\mathsf{int}}} \frac{s_N}{h} \gamma_\alpha [\![ \hat{c} ]\!] \cdot [\![ v_c ]\!] \; \mathrm{d}s \; , \tag{S21}$$

and hence arrive at the final weak form statement of Eq. (S11) which reads

$$\int_{\Omega_0} \frac{\partial}{\partial t} (\hat{J} \hat{c}) v_c \; \mathrm{d}\Omega_0 + \mathcal{D}(\hat{c}, v_c, \boldsymbol{\alpha}) - \int_{\Omega_0} \hat{J} R_c(t) v_c \; \mathrm{d}\Omega_0 = 0 \; . \tag{S22}$$

We note that an integral part of our model design is that continuity is only enforced on the internal edges. Thus jumps are possible only across the interface of the two subdomains i.e. across the membrane. In our case this means that the reactants in each cell cannot pass the cell-cell junction. This leads to a variety of possibilities in the treatment of multicellular systems. Cellular contractility in principle can be described by distinct reaction-diffusion systems in each cell. The RD-systems within the cells can then be coupled by appropriate mechano-chemical coupling terms to account for mechanosensing at the intercellular junction.

The weak form of Eq. (S11) finally reads

$$
\begin{aligned}
0 = &\int_{\Omega_0} \frac{\partial}{\partial t}(\hat{J}\hat{c}) v_c \; \mathrm{d}\Omega_0 + \int_{\Omega_0} \boldsymbol{\alpha}\hat{\nabla}\hat{c} \cdot \hat{\nabla} v_c \; \mathrm{d}\Omega_0 \\
&- \underbrace{\int_{\mathcal{F}_{\text{int}}} \{\boldsymbol{\alpha}\hat{\nabla}\hat{c}\}_\omega \cdot [\![ v_c ]\!] \; \mathrm{d}s}_{\text{consistency}} - \underbrace{\int_{\mathcal{F}_{\text{int}}} \{\boldsymbol{\alpha}\hat{\nabla} v_c\}_\omega \cdot [\![ \hat{c} ]\!] \; \mathrm{d}s}_{\text{symmetry}} + \underbrace{\int_{\mathcal{F}_{\text{int}}} \frac{s_N}{h}\gamma_\alpha [\![ \hat{c} ]\!] \cdot [\![ v_c ]\!] \; \mathrm{d}s}_{\text{penalty}} - \int_{\Omega_0} \hat{J} R_c(t) v_c \; \mathrm{d}\Omega_0 \, ,
\end{aligned}
$$

where $s_N$ denotes the Nitsche paramater, which must be chosen sufficiently large to ensure continuity across internal facets (7), and $h$ the average element diameter. The notation is illustrated in Fig. S1a.

#### Weak formulation for the elastic domain

To derive the weak formulation of

$$\nabla \cdot \boldsymbol{\sigma} = Y(\mathbf{x})\mathbf{u} \, , \tag{S23}$$

we multiply with a vector valued test function $\mathbf{v} \in \boldsymbol{\mathcal{V}}(\Omega_0)$ and integrate over the domain $\Omega_0$ of the undeformed configuration

$$\int_{\Omega_0} (\nabla \cdot \boldsymbol{\sigma}) \cdot \mathbf{v} \; \mathrm{d}\Omega_0 = \int_{\Omega_0} Y(\mathbf{x})\mathbf{u}(\mathbf{x}) \cdot \mathbf{v} \; \mathrm{d}\Omega_0 \, . \tag{S24}$$

The left hand side can be integrated using integration by parts i.e. using the following identity

$$\nabla \cdot (\boldsymbol{\sigma} \cdot \mathbf{v}) = (\nabla \cdot \boldsymbol{\sigma}) \cdot \mathbf{v} + \boldsymbol{\sigma} : \nabla \mathbf{v} \, . \tag{S25}$$

This allows to simplify Eq. (S24) to

$$\int_{\Omega_0} \boldsymbol{\sigma} : \nabla \mathbf{v} \; \mathrm{d}\Omega_0 - \int_{\Gamma} (\boldsymbol{\sigma} \cdot \mathbf{N}) \cdot \mathbf{v} \; \mathrm{d}s + \int_{\Omega_0} Y \mathbf{u} \cdot \mathbf{v} \; \mathrm{d}\Omega_0 = 0 \, . \tag{S26}$$

Here, $\boldsymbol{\sigma} \cdot \mathbf{N}$ is the traction vector at the boundary $\Gamma = \partial\Omega_0$ which is set to zero in case of stress free boundaries and hence, the final weak form statement reads

$$\int_{\Omega_0} \boldsymbol{\sigma} : \nabla \mathbf{v} \; \mathrm{d}\Omega_0 + \int_{\Omega_0} Y \mathbf{u} \cdot \mathbf{v} \; \mathrm{d}\Omega_0 = 0 \, . \tag{S27}$$

##### Time-discretisation

All time dependent quantities $Q(t)$ are discretized using an implicit (backward Euler) scheme at a given time $t^{(n+1)}$

$$\left(\frac{\mathrm{d}Q}{\mathrm{d}t}\right)^{(n+1)} \approx \frac{Q^{(n+1)} - Q^{(n)}}{\Delta t} \,. \tag{S28}$$

##### Mesh generation

The meshes for the three different systems were generated with Gmsh (9). For the cell chain and the tissue-like monolayer we made sure to create a mesh which respects the symmetry of the system. The meshes are shown in Fig. S1c.

#### Literature review RhoA-pathway

Having reviewed the relevant literature, we found that the model as presented by Kamps et al. (8) contains all important components which are necessary for a profound description of the RhoA pathway. In contrast to the RhoA-actomyosin system as introduced by Staddon et al. (10) it explicitly contains GEF as a downstream effector of RhoA, and thus provides an important interface for light-induced contraction as GEF activity can be controlled by optogenetic constructs like the CRY2/CIBN system. In combination with experimental measurements Kamps et al. (8) proposed a reaction scheme for the active reactants GEF (G), RhoA (R) and myosin (M) which reads

$$\frac{\mathrm{d}G}{\mathrm{d}t} = k_3 R (G_T - G) - k_4 G M \tag{S29}$$

$$\frac{\mathrm{d}R}{\mathrm{d}t} = \frac{k_1 G (R_T - R)}{K_{m1} + R_T - R} - k_2 \frac{R}{K_{m2} + R} \tag{S30}$$

$$\frac{\mathrm{d}M}{\mathrm{d}t} = \frac{k_5 R (M_T - M)}{K_{m5} + M_T - M} - k_6 \frac{M}{K_{m6} + M} \,. \tag{S31}$$

$G_T, R_T$ and $M_T$ denote the total concentrations of the species which the authors assume to be constant. The rate constants are denoted by $k_i$ and the Michaelis-Menten constants are given by the $K_{mi}$.

The membrane and cytosol associated species represent the active and passive states, respectively. This terminology stems from experimental studies which show that the active forms of RhoA and myosin are predominantly found in the vicinity of the plasma membrane and the submembraneous actin cortex. In contrast, the inactive forms are associated with the cytosol (11, 12). The RhoA protein for example exhibits a lipophilic end which enables it to bind to lipid membranes (13). However, so-called guanosine dissociation inhibitors (GDIs) may bind to Rho-GDPs, not only keeping them in a permanently inactive state but also preventing its membrane localization by shielding the hydrophilic end and additionally making it soluble in the cytoplasm (14). The reaction scheme also highlights the two feedback loops which are important to describe the excitable and oscillatory dynamics that are observed in experiments (see Fig. S1 d). The positive feedback loop stems from the observation that RhoA activity at the membrane further induces GEF membrane recruitment. Due to Rho activation by GEFs, this closes a positive feedback loop (8). The negative feedback loop can be traced back to the ability of myosin to inhibit the nucleotide exchange activity of GEFs by binding to their Dbl-homology domain (DH) (15). Essentially, the authors could identify the total concentration of active GEF as the main bifurcation parameter for the switch from stable to oscillatory states at intermediate GEF concentrations. In experiments, they vary this bifurcation parameter by treating cells with nocodazole, which leads to depolymerization of microtubules from which GEFs are then released. The crossover from stable to osciallatory dynamics happens as a function of the total GEF concentration.

### Parametrization

#### Non-linear RhoA pathway

For the simulations with the non-linear Rho pathway we follow Kamps et al. (8). In Eqs. (S29) to (S31) we treat the inactive species separately with $c_i = c_T - c$. The inactive species diffuse faster than the active species $D_{c,i} > D_c$. The reaction kinetics are given by $R_{c_i} = -R_c$. Hence, for the non-linear system we have a total of six coupled reaction diffusion equations coupled to the PDE describing the cell layer. Fig. S1d shows the schematic of the coupled system of PDE's. The reaction diffusion system alone exhibits instabilities that lead to the emergence of traveling wave peaks. To induce an instability we impose initial conditions on the active species by adding small random fluctuations of the form

$$c_0(\mathbf{x}) = \bar{c}_0 + \delta c_0(0.5 - \mathcal{U}_{[0,1](\mathbf{x})}) \,, \tag{S32}$$

where $\mathcal{U}_{[0,1](\mathbf{x})}$ is the probability density function of the continuous uniform distribution, $\bar{c}_0$ the homogeneous concentration field and $\delta c_0$ a small flucutation. For the inactive species we set $c_{i,0}(\mathbf{x}) = c_T - c_0(\mathbf{x})$. Fluctuations are $\delta G_0 = 0.05$, $\delta R_0 = 0.01$ and $\delta M_0 = 0$. All other relevant parameters can be found in Tab. S1 or in the supplemental information of (8).

#### Linearized RhoA pathway

For the parametrization of the proposed linear signaling cascade we rely on the order of magnitudes found in the respective literature. Within the limits of our simplified model the total concentrations of RhoA and myosin are irrelevant since they do not explicitly enter the reaction kinetics in the weakly activated regime. The parameters are chosen such that the steady state concentrations of RhoA and myosin are roughly $10\,\%$ of the total concentration (10, 16). Further, the reaction rates are chosen such that the time course of the myosin concentration approximates the typical time course of actively generated stresses during optogenetic activation (8, 10, 17–19). The time course of the input signal was adapted to the measured CRY2 membrane recruitment and is described by a relaxation time of $\approx 10^2\,\mathrm{s}$ $(\lambda \approx 0.01\,\mathrm{s}^{-1})$(20). The reference values from (8) were estimated as follows. The second term in Eq. (S30) can be approximated in a weakly activated regime with $K_{m2} \gg R$ as $k_2 R/(K_{m2} + R) \approx k_2 R/K_{m2} \equiv b \approx 2\,\mathrm{s}^{-1}$. The same argument applied to the second term in

Eq. (S31) gives $k_6 M/(K_{m6}+M) \approx k_6 M/K_{m6} \equiv s \approx 0.0051\,\mathrm{s}^{-1}$. $k$ can be estimated from the first term in Eq. (S31) by $k_5/(K_{m5}+M_T) = \tilde{k} = 0.004\,65\,\mathrm{s}^{-1}$ from which $k = \tilde{k} R_T = 0.002\,\mathrm{s}^{-1}$ follows. All relevant parameters are summarized in Tab. S2.

##### Elastic layer

For the parametrisation of the cell and the substrate we follow the typical orders of magnitude. Cell and substrate have a Young's modulus $E$ in the range of several $\mathrm{kPa}$. For simplicity we choose $E_c \approx E_s$, which also reflects that cells typically adapt to the stiffness of their environment. The viscoelastic time scale $\tau_c$ is a free parameter in the simulations and hence the viscosity of the cell layer is defined by $\eta_c = \tau_c E_c$. The spring stiffness density is calculated by $Y_s = \pi E_s/L_c$ (21), where $L_c$ is the lateral extent of a cell. All other relevant parameters are summarized in Tables S3 to S5.

#### Analytical solution of the weakly activated signaling cascade

Beguerisse-Díaz et al. (22) provide a variety of analytical solutions to weakly activated signalling cascades triggered by different input signals i.e. time course of the stimulus such as step-function, Gaussian or, as in our case, an exponential decreasing perturbation. The signaling species $x_1^*$ is activated by an external stimulus, which in turn activates species $x_2^*$, and so on. In a weakly activated regime and for $x_i^*(0) = 0$ (which is the initial condition for the perturbations $\delta r$ and $\delta m$) the output function of species $x_n^*$ is given by

$$x_n^*(t) = \left(\prod_{i=1}^{n} \alpha_i\right) \sum_{i=1}^{n} \left(\prod_{q=1, q\neq i}^{n} (\beta_i - \beta_q)^{-1}\right) \times \int_0^t e^{-\beta_i (t-\tau)} A(\tau)\ \mathrm{d}\tau \,. \tag{S33}$$

Here $\alpha_i$ and $\beta_i$ denote the activation and deactivation rates of each species $i$, respectively, and $A(t)$ is a stimulus applied to the first species. Applied to the system of equations

$$\frac{\mathrm{d}\delta r}{\mathrm{d}t} = a\delta g(t) - b\delta r\,, \quad \frac{\mathrm{d}\delta m}{\mathrm{d}t} = k\delta r - s\delta m\,, \tag{S34}$$

together with $A(\tau) = \delta g(\tau) = g_a e^{-\lambda \tau}$ (as in the main text) this leads to

$$\delta\tilde{r}(t) = \frac{\delta r}{r_{\mathsf{ss}}} = \frac{b\alpha}{b-\lambda}\left(e^{-\lambda t} - e^{-bt}\right)\,, \tag{S35}$$

$$\delta\tilde{m}(t) = \frac{\delta m}{m_{\mathsf{ss}}} = b\alpha s\left(\frac{e^{-\lambda t} - e^{-bt}}{(b-s)(b-\lambda)} - \frac{e^{-\lambda t} - e^{-st}}{(b-s)(s-\lambda)}\right)\,. \tag{S36}$$

### Description of movies

**Movie S1:** Strain-dependent feedback in a cell doublet with strong coupling. Activation of the left cell leads to substantial contraction in the right cell. Although optogenetic stimulation is only applied to the left cell, the whole doublet contracts as a unit i.e. symmetrically. Parameters used as listed in Tab. S2 and Tab. S3 with $a_{\delta\tilde{q}} = 100$ and $\tau_c = 10\text{s}$.

**Movie S2:** Strain-dependent feedback in a cell doublet with weak coupling. Activation of the left cell leads to weak contraction in the right cell. This leads to an overall asymmetric shape deformation of the whole doublet in which the right cell gets pulled to the left. Parameters used as listed in Tab. S2 and Tab. S3 with $a_{\delta\tilde{q}} = 0.1$ and $\tau_c = 10\text{s}$.

**Movie S3:** Propagation of a contraction wave through a cell chain visualized by the deformation field (color code: red color corresponds to displacement to the right, blue to the left). Optogenetic activation of the left cell leads to a contraction. The coupling to the neighboring cell induces a contraction which propagates from the left end of the cell chain to the right end. Parameters used as listed in Tab. S2 and Tab. S4. Coupling and viscoelastic time scale where chosen from the transmissive regime. Red lines indicate initial cell-cell boundary positions.

**Movie S4:** Propagation of contraction wave through a tissue-like monolayer comprised of 28 cells. Here, the contractility is controlled by the non-linear Rho-pathway which in itself exhibits wave-like instabilities. The parameters are chosen according to Tab. S1 and Tab. S5. At early times, an instability is triggered in the centered cell on the left by adding small random fluctuations on the GEF and Rho component. These random fluctuations lead to several concentrated traveling wave peaks which in turn trigger GEF activation in adjacent cells through the strain-dependent coupling. Thus, a contraction wave spreads through the whole tissue activating all cells and triggering the wave-like instability. The emerging activation pattern represents the up-down symmetry of the tissue. Cells at the free edge (edge with no cell-cell boundary) strongly deform when a contraction peak gets close to the free cell edge.

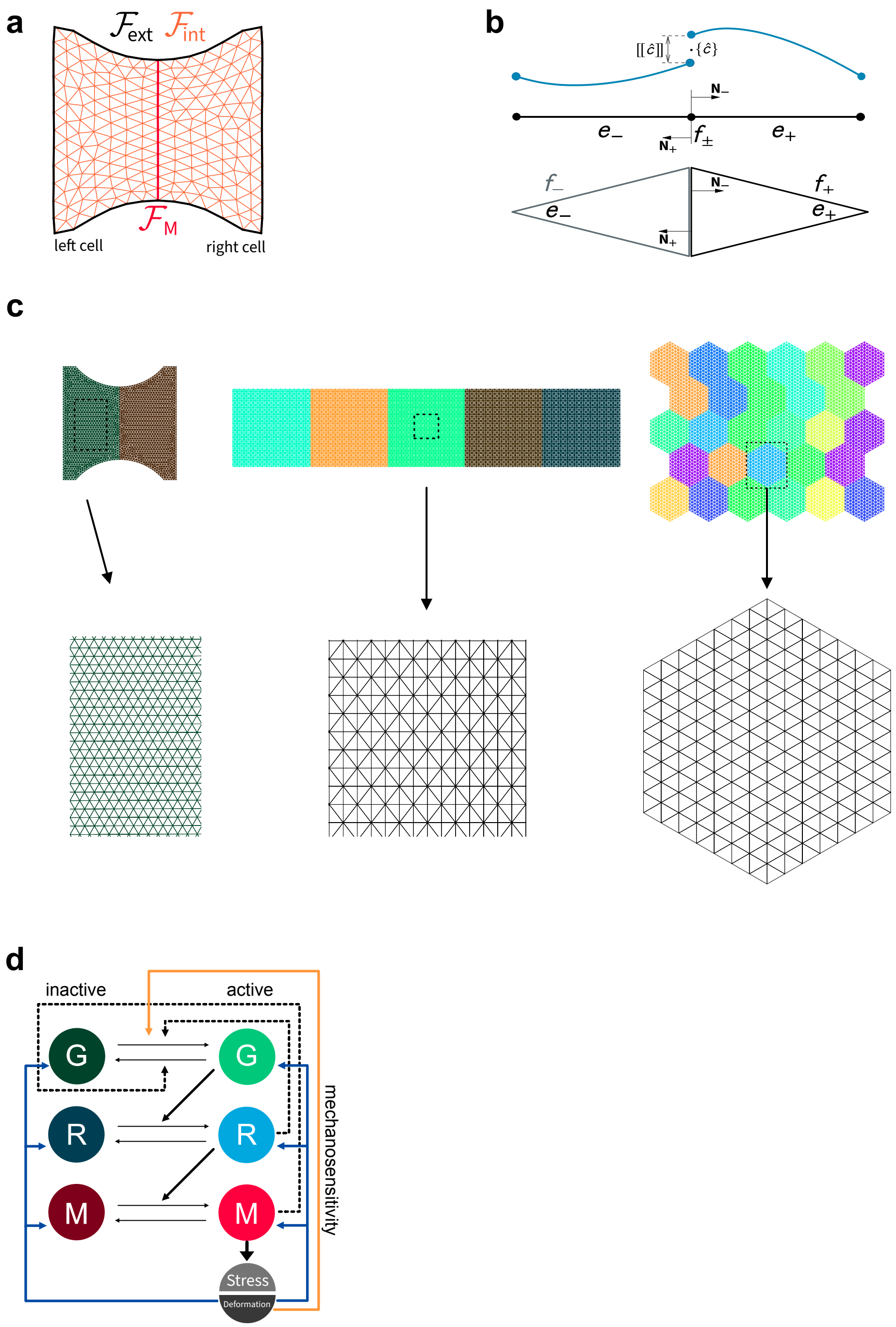


Figure S1: Panel (a) and (b) schematically explains the notation used in the derivation of the discontinuous Galerkin finite element method. Panel (c) depicts the meshes for the three cell systems. Colors highlight different cells. For cell chain and tissue-like monolayer we enforced meshes to respect the symmetry of the system's geometry. Panel (d) shows the coupled system of the non-linear RhoA pathway by Kamps et al. (8) coupled to the mechanics of the elastic layer. Mechanical feedbacks represented by yellow and blue arrows.

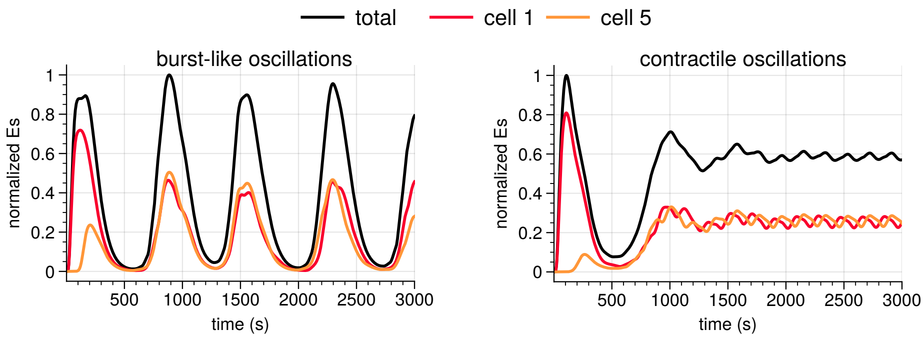


Figure S2: Different oscillations observed in a cell chain coupled to the linearized RhoA pathway. Burst-like oscillations (left) correspond to large variations in strain energy while contractile oscillations (right) corresponds to states in which the whole cell chain remains on average in a state of non-vanishing strain energy.

| Abbreviation | Value |
|---|---|
| | GEF |
| $G_T$ | $0.2\mathrm{M}$ |
| $k_3$ | $1.19\mathrm{M}^{-1}\mathrm{s}^{-1}$ |
| $k_4$ | $3.98\mathrm{M}^{-1}\mathrm{s}^{-1}$ |
| $D_G$ | $0.3\,\mu\mathrm{m}^2\,\mathrm{s}^{-1}$ |
| $D_{G_i}$ | $9.28\,\mu\mathrm{m}^2\,\mathrm{s}^{-1}$ |
| | RhoA |
| $R_T$ | $0.443\mathrm{M}$ |
| $K_{m1}$ | $2.42\mathrm{M}$ |
| $K_{m2}$ | $0.0745\mathrm{M}$ |
| $k_1$ | $(3.88K_{m1})\mathrm{s}^{-1}$ |
| $k_2$ | $(2.04K_{m2})\mathrm{Ms}^{-1}$ |
| $D_R$ | $0.3\,\mu\mathrm{m}^2\,\mathrm{s}^{-1}$ |
| $D_{R_i}$ | $9.28\,\mu\mathrm{m}^2\,\mathrm{s}^{-1}$ |
| | Myosin |
| $M_T$ | $1.24\mathrm{M}$ |
| $K_{m5}$ | $0.014\mathrm{M}$ |
| $K_{m6}$ | $0.784\mathrm{M}$ |
| $k_5$ | $(0.417K_{m5})\mathrm{s}^{-1}$ |
| $k_6$ | $(0.00509K_{m9})\mathrm{Ms}^{-1}$ |
| $D_M$ | $0.03\,\mu\mathrm{m}^2\,\mathrm{s}^{-1}$ |
| $D_{M_i}$ | $0.9\,\mu\mathrm{m}^2\,\mathrm{s}^{-1}$ |

Table S1: Parameter values for the non-linear Rho pathway with fast and slow diffusing species according to (8). Here, we use $\mathrm{M}$ to indicate units of concentration which corresponds to $10^6$ molecules per cell (as given in the SI of (8)).

| Abbreviation | Used value | Ref. value | Reference |
|---|---|---|---|
| $\lambda$ | $0.01\,\mathrm{s}^{-1}$ | $0.008\,\mathrm{s}^{-1} - 0.018\,\mathrm{s}^{-1}$ | (20) |
| $\alpha$ | $100\,\%$ | $20\,\% - 130\,\%$ | (20) |
| $b$ | $0.0165\,\mathrm{s}^{-1}$ | $2\,\mathrm{s}^{-1}$ | (8) |
| $k$ | $0.1\,\mathrm{s}^{-1}$ | $0.1408\,\mathrm{s}^{-1}$ | (10) |
| | | $0.002\,\mathrm{s}^{-1}$ | (8) |
| $s$ | $0.083\,\mathrm{s}^{-1}$ | $0.0051\,\mathrm{s}^{-1}$ | (8) |
| | | $0.082\,\mathrm{s}^{-1}$ | (10) |
| $m_{\mathrm{ss}}$ | $0.1$ | $0.1 - 0.8$ | (10) |
| | | | (16) |
| $D_R, D_G$ | $0.3\,\mathrm{\mu m}^2\,\mathrm{s}^{-1}$ | $0.28\,\mathrm{\mu m}^2\,\mathrm{s}^{-1}$ | (8) |
| | | $0.1\,\mathrm{\mu m}^2\,\mathrm{s}^{-1}$ | (23) |
| $D_M$ | $0.03\,\mathrm{\mu m}^2\,\mathrm{s}^{-1}$ | $0.03\,\mathrm{\mu m}^2\,\mathrm{s}^{-1}$ | (8) |
| | | $0.01\,\mathrm{\mu m}^2\,\mathrm{s}^{-1}$ | (23) |
| | Deduced | | |
| $r_{\mathrm{ss}}$ | $0.083$ | | |
| a | $0.0014\,\mathrm{s}^{-1}$ | $< 0.002\,\mathrm{s}^{-1}$ | (10) |

Table S2: Parameter values for the linearized RhoA signaling cascade. We set most of the parameters in accordance with the reported ranges. The parameters taken from (8) where obtained by taking the corresponding activation and deactivation rates in a weakly activated regime for which the Michaelis-Menten terms can be linearly approximated. However, their model does not provide a basal activation rate. The reported rate constants as stated in the work of Staddon et al. (10) were deduced from Michaux et al. (24). The parameters $r_{\mathrm{ss}}$ and $a$ are not independent and consequently deduced from the fixed parameters.

| Abbreviation | Value |
|---|---|
| Cell parameters | |
| Young's Modulus $E_c$ | $5 \cdot 10^3 \mathrm{Pa}$ |
| Poisson's ratio $\nu_c$ | $0.5$ |
| Cell height $h_c$ | $1 \cdot 10^{-6} \mathrm{m}$ |
| Lateral extent $L_c$ | $45 \cdot 10^{-6} \mathrm{m}$ |
| Two-dimensional active stress $\sigma_0$ | $2 \cdot 10^{-3} \mathrm{N\,m^{-1}}$ |
| Substrate parameters | |
| Young's Modulus $E_s$ | $5 \cdot 10^3 \mathrm{Pa}$ |
| Poisson's ratio $\nu_s$ | $0.5$ |

Table S3: Parameter values for the simulation of the cell doublet on an H-pattern

| Abbreviation | Value |
|---|---|
| Cell parameters | |
| Young's Modulus $E_c$ | $5 \cdot 10^3 \mathrm{Pa}$ |
| Poisson's ratio $\nu_c$ | $0.5$ |
| Cell height $h_c$ | $1 \cdot 10^{-6} \mathrm{m}$ |
| Lateral extent $L_c$ | $40 \cdot 10^{-6} \mathrm{m}$ |
| Two-dimensional active stress $\sigma_0$ | $2 \cdot 10^{-3} \mathrm{N\,m^{-1}}$ |
| Substrate parameters | |
| Young's Modulus $E_s$ | $2.5 \cdot 10^3 \mathrm{Pa}$ |
| Poisson's ratio $\nu_s$ | $0.5$ |

Table S4: Parameter values for the simulation of the cell chain with continuous adhesion and unidirectional contraction.

| Abbreviation | Value |
| --- | --- |
| Cell parameters | |
| Young's Modulus $E_c$ | $2.5 \cdot 10^3 \mathrm{Pa}$ |
| Poisson's ratio $\nu_c$ | $0.5$ |
| Cell height $h_c$ | $1 \cdot 10^{-6} \mathrm{m}$ |
| Lateral extent $L_c$ | $40 \cdot 10^{-6} \mathrm{m}$ |
| Two-dimensional active stress $\sigma_0$ | $2.5 \cdot 10^{-3} \mathrm{N\,m^{-1}}$ |
| Substrate parameters | |
| Young's Modulus $E_s$ | $1 \cdot 10^3 \mathrm{Pa}$ |
| Poisson's ratio $\nu_s$ | $0.5$ |

Table S5: Parameter values for the simulation of the tissue-like monolayer with non-linear Rho-pathway. Here, we chose a smaller Young's modulus for the cell, a softer substrate and a slightly larger value for active stress in order to allow for more visible deformations.

### Example for python FEM-code for photoactivation of a cell doublet on H-pattern

Our model was implemented in the FEM-framework FEniCS (25). This is an example code for the photoactivation of a cell doublet with mechanochemical feedback.

```python
# Import required libraries for finite element analysis and data processing
from dolfin import *           # FEniCS/DOLFIN for finite element computations
import numpy as np             # Numerical computations
import random                  # Random number generation
import copy                    # Deep copying of objects
import matplotlib.pyplot as plt  # Plotting utilities
from ufl import tanh          # Hyperbolic tangent function
from numpy import savetxt     # Array saving utility
import pandas as pd           # Data manipulation and analysis
import os                     # Operating system interface
import csv                    # CSV file operations

''' Finite-Element-Simulation of a coupled system of reaction-diffusion equations and 2D
    viscoelasticity

    This code implements a coupled mechanochemical model for cell mechanics using FEM.
    The model combines:
    1. Reaction-diffusion equations for biochemical species
    2. 2D viscoelastic mechanics for cell deformation
    3. Mechanochemical feedback between strain and signaling

    Copyright: Dennis Woerthmueller
    Date: February 14, 2024
'''

# Create data directory for simulation outputs
path_to_rawData = 'Data/'
if not os.path.exists(path_to_rawData):
    os.makedirs(path_to_rawData)

# Load simulation parameters from CSV file
with open('inputParams.csv', newline = '') as file:
    reader = csv.reader(file, quoting = csv.QUOTE_NONNUMERIC,
                        delimiter = ' ')
    rows = list(reader)
    keys = rows[0]      # Parameter names
    values = rows[1]    # Parameter values

# Create parameter dictionary from keys and values
key_value_pairs = zip(keys, values)
p = dict(key_value_pairs)

def calculateStrainEnergy(u, kN, dx):
    """
    Calculate the elastic strain energy in the substrate.

    Args:
        u (Function): Displacement field
        kN (Expression): Spring constant field
        dx (Measure): Integration measure

    Returns:
        float: Total strain energy
    """
    return assemble(0.5*kN*inner(u,u)*dx)

class KNExpression(UserExpression):
```

```python
    """
    Define position-dependent spring constants for an H-shaped micropattern.

    The pattern consists of two vertical arms connected by a horizontal crossbar.
    Spring constants are non-zero only within the pattern.
    """
    def __init__(self,Y, armWidth, degree=2):
        print("BIS HIER")
        super().__init__()
        self.armWidth = armWidth  # Width of pattern arms
        self.Y = Y                # Young's modulus / spring constant

    def eval(self, value, x):
        """
        Evaluate spring constant at given position.

        Args:
            value: Output value (modified in-place)
            x: Spatial coordinates
        """
        d = 1  # Domain size
        # Set spring constant Y in arms and crossbar, 0 elsewhere
        if (x[0] <= -(d/2)+self.armWidth or x[0] >= (d/2)-self.armWidth or
            between(x[1],(-self.armWidth/2,self.armWidth/2))):
            value[0] = self.Y
        else:
            value[0] = 0.0

def normalize_solution(U, max):
    """
    Normalize solution vector by dividing by maximum value.

    Args:
        U (Function): Solution vector to normalize
        max (float): Maximum value to normalize by

    Returns:
        Function: Normalized solution vector
    """
    U_array = U.vector().get_local()
    U_array /= max
    U.vector()[:] = U_array
    return U

def eps(v):
    """
    Calculate strain tensor from displacement field.

    Args:
        v (Function): Displacement field

    Returns:
        Tensor: Symmetric gradient (strain tensor)
    """
    return sym(grad(v))

class selectedSubdomain(UserExpression):
    """
    Mark specific subdomains with different values.
    Used to identify and assign properties to different regions.
    """
    def __init__(self, subdomains, val_inside, val_outside, subdomain_id, **kwargs):
        super().__init__()
        self.subdomains = subdomains        # Subdomain markers
        self.val_inside = val_inside        # Value inside selected subdomain
        self.val_outside = val_outside      # Value outside selected subdomain
```

```python
        self.subdomain_id = subdomain_id    # ID of subdomain to mark

    def eval_cell(self, values, x, cell):
        """
        Evaluate marker value for each cell.

        Args:
            values: Output value (modified in-place)
            x: Spatial coordinates
            cell: Current cell
        """
        if self.subdomains[cell.index] == self.subdomain_id:
            values[0] = self.val_inside
        else:
            values[0] = self.val_outside

class SquareCompartmentDoublet(UserExpression):
    """
    Define square compartment for cell doublet simulation.
    Used to create initial conditions and activation patterns.
    """
    def __init__(self,A,d, degree=0):
        super().__init__()
        self.A = A  # Amplitude
        self.d = d  # Distance/size parameter

    def eval(self, value, x):
        """
        Evaluate compartment value at given position.

        Args:
            value: Output value (modified in-place)
            x: Spatial coordinates
        """
        if (x[0] <= -self.d):
            value[0] = self.A
        else:
            value[0] = 0.0

def get_boundary_of_deformed_mesh(u, geo_file_name):
    """
    Extract boundary coordinates of deformed mesh.
    Used for tracking boundary deformation over time.

    Args:
        u (Function): Displacement field
        geo_file_name (str): Base name of geometry files

    Returns:
        numpy.array: Sorted boundary coordinates
    """
    # Load mesh and boundary definitions
    dummy_mesh = Mesh("%s.xml"%(geo_file_name))
    boundaries = MeshFunction("size_t", dummy_mesh, "%s_facet_region.xml"%(geo_file_name))
    subdomains = MeshFunction("size_t", dummy_mesh, "%s_physical_region.xml"%(geo_file_name))

    # Apply displacement to mesh
    ALE.move(dummy_mesh, u)
    V_mesh = FunctionSpace(dummy_mesh, "CG", 1)
    v2d = vertex_to_dof_map(V_mesh)

    # Extract boundary vertices
    dofs = []
    for facet in facets(dummy_mesh):
        if boundaries[facet.index()] == 2:  # Top boundary
            vertices = facet.entities(0)
```

```
            for vertex in vertices:
                dofs.append(v2d[vertex])

    # Sort and return boundary coordinates
    unique_dofs = np.array(list(set(dofs)), dtype=np.int32)
    boundary_coords = V_mesh.tabulate_dof_coordinates()[unique_dofs]
    col = 0
    boundary_coords_sorted = boundary_coords[np.argsort(boundary_coords[:,col])]
    return boundary_coords_sorted

def DGWeakFormRD(c, cn, vc, Dc, u, J, J_n, F, n, dx, dS, dSM, dt, cellularisation=True):
    """
    Construct weak form for reaction-diffusion equations using Discontinuous Galerkin method.

    Args:
        c (Function): Current concentration
        cn (Function): Previous concentration
        vc (TestFunction): Test function
        Dc (float): Diffusion coefficient
        u (Function): Displacement field
        J (Expression): Current Jacobian
        J_n (Expression): Previous Jacobian
        F (Expression): Deformation gradient
        n (Expression): Normal vector
        dx (Measure): Volume measure
        dS (Measure): Interior facet measure
        dSM (Measure): Interface measure
        dt (float): Time step
        cellularisation (bool): Whether to include cell interface terms

    Returns:
        Form: Complete weak form for reaction-diffusion equation
    """
    # Calculate geometric quantities
    F_inv = inv(F)
    F_inv_T = inv(F).T
    I = Identity(2)
    eps = sym(grad(u))

    # Numerical parameters
    h = 0.1   # Mesh size parameter
    sN = 50   # Penalty parameter

    # Modified diffusion tensor including geometric factors
    alph = J*(I-2*eps)*Dc

    # Calculate interface terms for DG formulation
    delt_p = dot(n('+'), alph('+')*n('+'))
    delt_m = dot(n('-'), alph('-')*n('-'))

    # Weights for averaging
    w_pos = delt_m/(delt_p+delt_m)
    w_neg = delt_p/(delt_p+delt_m)

    # Average terms for concentration gradients
    grad_c_avg_term = w_pos*alph('+')*grad(c)('+')+w_neg*alph('-')*grad(c)('-')
    grad_vc_avg_term = w_pos*alph('+')*grad(vc)('+')+w_neg*alph('-')*grad(vc)('-')

    # Interface penalty parameter
    gamma = 2*delt_p*delt_m/(delt_p+delt_m)

    # Time derivative terms
    time_deriv = J*(c-cn)/dt*vc*dx+(J-J_n)/dt*c*vc*dx

    # Standard diffusion term
    standard = dot(alph * grad(c), grad(vc)) * dx
```

```

    # Interface terms based on cellularisation flag
    if cellularisation:
        # Terms for internal cell boundaries
        consistency = -dot(jump(vc,n),grad_c_avg_term)*dS + dot(jump(vc,n),grad_c_avg_term)*dSM
        symmetry = - dot(grad_vc_avg_term, jump(c,n))*dS + dot(grad_vc_avg_term,jump(c,n))*dSM
        penalty = sN/h*gamma*dot(jump(vc, n), jump(c,n))* dS - sN/h*gamma*dot(jump(vc,n), jump(c, n))*dSM
    else:
        # Terms without internal boundaries
        consistency = -dot(jump(vc,n),grad_c_avg_term)*dS
        symmetry = - dot(grad_vc_avg_term, jump(c,n))*dS
        penalty = sN/h*gamma*dot(jump(vc, n), jump(c,n))* dS

    # Combine all terms
    weakForm = time_deriv + standard + consistency + symmetry + penalty

    return weakForm

# Output file names
name = 'simulation_output'           # Name of the output xdmf-file
geo_file_name = 'cell_doublet_shape_nonDim'  # Name of gmsh .geo-file

def simulation():
    """
    Main simulation function implementing a mechanochemical feedback model.

    The simulation couples three main components:
    1. Reaction-diffusion system for GEF-RhoA-Myosin signaling
    2. Viscoelastic mechanics for cell deformation
    3. Mechanochemical feedback through strain

    Uses mixed finite elements: DG for concentrations, CG for displacement.
    """
    # ------------------------------------------------------------------------
    # Initialize mesh and finite element structures
    # ------------------------------------------------------------------------
    # Convert mesh from Gmsh format to FEniCS XML format
    if not os.path.exists('%s.xml'%(geo_file_name)):
        os.system('dolfin-convert %s.msh %s.xml'%(geo_file_name,geo_file_name))

    # Load mesh and domain definitions
    mesh = Mesh("%s.xml"%(geo_file_name))
    boundaries = MeshFunction("size_t", mesh, "%s_facet_region.xml"%(geo_file_name))
    subdomains = MeshFunction("size_t", mesh, "%s_physical_region.xml"%(geo_file_name))

    # Save domain definitions for visualization
    file_results = XDMFFile("subdomains.xdmf")
    file_results.write(subdomains)
    file_results = XDMFFile("boundaries.xdmf")
    file_results.write(boundaries)

    # Define measures for integration
    dx = Measure('dx', domain=mesh, subdomain_data=subdomains)    # Volume measure
    dS_all = Measure('dS', subdomain_data=boundaries)            # Surface measure
    dSM = dS_all(1)                                             # No-flux interface measure

    # Initialize domain markers
    cell1 = selectedSubdomain(subdomains, 1, 0, subdomain_id = 1, degree=0)       # Active cell
    non_opto_cell = selectedSubdomain(subdomains, 0, 1, subdomain_id = 1, degree=0) # Non-photoactivated cell
    n = FacetNormal(mesh)                                                          # Normal vector field

    # Define finite elements for mixed formulation
    P1 = FiniteElement('DG', triangle, 1)           # DG elements for concentrations
```

```
    P2 = VectorElement('CG', triangle, 1)             # CG elements for displacement
    element = MixedElement([P1, P1, P1,P2])           # Combined element
    V = FunctionSpace(mesh, element)                  # Mixed function space

    # Initialize function spaces for output fields
    # DG spaces for scalar fields
    dFE_DG0 = FiniteElement("DG", mesh.ufl_cell(), 0)
    dFE_DG1 = FiniteElement("DG", mesh.ufl_cell(), 1)
    dFE_CG1 = FiniteElement("CG", mesh.ufl_cell(), 1)
    dFE_CG2 = FiniteElement("CG", mesh.ufl_cell(), 2)

    # Tensor spaces for stress/strain fields
    TensorSpace_DG0 = TensorFunctionSpace(mesh, "DG", 0)
    TensorSpace_DG1 = TensorFunctionSpace(mesh, "DG", 1)
    TensorSpace_CG1 = TensorFunctionSpace(mesh, "CG", 1)
    TensorSpace_CG2 = TensorFunctionSpace(mesh, "CG", 2)

    # Create function spaces
    W_DG0 = TensorSpace_DG0     # DG0 tensor space
    W_DG1 = TensorSpace_DG1     # DG1 tensor space
    W_CG1 = TensorSpace_CG1     # CG1 tensor space
    W_CG2 = TensorSpace_CG2     # CG2 tensor space

    # Scalar function spaces
    K_DG0 = FunctionSpace(mesh, dFE_DG0)
    K_DG1 = FunctionSpace(mesh, dFE_DG1)
    K_CG1 = FunctionSpace(mesh, dFE_CG1)
    K_CG2 = FunctionSpace(mesh, dFE_CG2)

    # Vector function space for displacement
    V_CG1 = VectorFunctionSpace(mesh,"CG",1)

    # Initialize output functions
    # Vector fields (displacement and traction)
    disp = Function(V_CG1, name='Displacement')
    TractionF = Function(V_CG1, name='Traction')

    # Scalar fields for molecular species
    GEF = Function(K_DG0, name='GEF')                      # GEF concentration
    RhoA = Function(K_DG0, name='RhoA')                    # RhoA concentration
    Myosin = Function(K_DG0, name='Myosin')              # Myosin concentration
    GEF_inactive = Function(K_DG0, name='GEF inactive')
    RhoA_inactive = Function(K_DG0, name='RhoA inactive')
    Myosin_inactive = Function(K_DG0, name='Myosin inactive')

    # Scalar fields for mechanics
    activatedCell = Function(K_DG0, name='Activated Cell')
    nonOptoCells = Function(K_DG0, name='Non-opto Cells')
    pattern = Function(K_DG0, name='Micropattern')
    Jacobian = Function(K_DG0, name='detF')
    JacobianPositive = Function(K_DG0, name='detF +')
    feedbackPositive = Function(K_DG0, name='feedback')
    hypTangentPositive = Function(K_DG0, name='tanh')
    traceGreenLagrange = Function(K_DG0, name='trE')
    traceGreenLagrangePositive = Function(K_DG0, name='trE +')
    detCauchyStressPositive = Function(K_DG0, name='det(CS) +')

    # Tensor fields for stress and strain
    Cauchystress = Function(W_DG0, name='Cauchy Stress')
    Cauchystress_passive = Function(W_DG0, name='Passive Cauchy Stress')
    Pstress = Function(W_DG0, name='Piola1 Stress')
    Pstress_passive = Function(W_DG0, name='Passive Piola1 Stress')
    activeStress = Function(W_DG0, name='Active Stress')
    strainGreenLagrange = Function(W_DG0, name='E (GL Strain)')
    defGrad_save = Function(W_DG0, name='Deformation Gradient Tensor')
    CauchyGreenInverse_save = Function(W_DG0, name='Inverse Cauchy Green')
```

```
    # Diffusion tensors
    diffTensor_G = Function(W_DG0, name='alpha_G')
    diffTensor_R = Function(W_DG0, name='alpha_R')
    diffTensor_M = Function(W_DG0, name='alpha_M')

    # Initialize output file
    xdmf_file = XDMFFile(path_to_rawData+"%s.xdmf"%(name))
    xdmf_file.parameters["flush_output"] = True
    xdmf_file.parameters["functions_share_mesh"] = True

    # Define spring constant field for substrate
    kN = KNExpression(p['Ys_N'], p['armWidth_N'], degree=2)

    # Set time discretization parameters
    DT = 0.5                   # Time step size
    dt = Constant(DT)          # FEniCS constant for time step
    t = DT                     # Current time
    T = 500                    # End time
    t_opto = 5                  # photoactivation

    # Initialize variational problem components
    dU = TrialFunction(V)      # Trial function
    U_tot = Function(V)        # Current solution
    U_tot_n = Function(V)      # Previous solution
    vG, vR, vM, vu = TestFunctions(V)  # Test functions

    # Additional functions for solution storage
    U_tot_save = Function(V)
    u_save = Function(V_CG1)
    U_tot_save_n = Function(V)

    # Set initial conditions
    u0 = Constant((0.0,0.0))       # Zero displacement
    random.seed()                   # Set random seed
    G_0 = Constant(0)              # Initial GEF concentration
    R_0 = Constant(0)              # Initial RhoA concentration
    M_0 = Constant(0)              # Initial Myosin concentration

    # Project initial conditions to appropriate function spaces
    uG_n = project(G_0, V.sub(0).collapse())
    uR_n = project(R_0, V.sub(1).collapse())
    uM_n = project(M_0, V.sub(2).collapse())
    u_n = project(u0, V.sub(3).collapse())

    # Assign initial conditions to solution vector
    assign(U_tot_n, [uG_n,uR_n,uM_n,u_n])

    # Split solution for component access
    uG,uR, uM, u = split(U_tot)            # Current solution
    uG_n, uR_n, uM_n, u_n = split(U_tot_n)  # Previous solution

    # Initialize arrays for storing results
    time_array = []
    strainEnergyLeft_vs_time = []
    strainEnergyRight_vs_time = []
    strainEnergyTotal_vs_time = []
    boundary_curve_vs_time = []



    # Main time stepping loop
    while t <= T:
        print("TIME = ", t)

        # -------------------------------------------------------------------------------------
```

```
        # Set up tensors for continuum description of the cell layer
        # -------------------------------------------------------------------------------------
        # Define fundamental geometric tensors
        I = Identity(2)                    # Identity tensor
        F = I + grad(u)                    # Deformation gradient tensor
        J_n = 1+tr(grad(u_n))             # Previous Jacobian (volume change)
        J = 1+tr(grad(u))                 # Current Jacobian
        F_transpose = F.T                  # Transpose of deformation gradient
        F_inv = I - grad(u)                # Approximate inverse of deformation gradient
        C = I - 2*sym(grad(u))            # Right Cauchy-Green tensor
        C1 = inv(C)                        # Inverse of Cauchy-Green tensor

        # Calculate strain measures
        eps_n = sym(grad(u_n))            # Previous strain tensor
        eps = sym(grad(u))                # Current strain tensor
        trace_n = tr(eps_n)               # Previous volumetric strain
        trace = tr(eps)                   # Current volumetric strain

        # -------------------------------------------------------------------------------------
        # Define active stresses from myosin activity
        # -------------------------------------------------------------------------------------
        # Calculate myosin-dependent active stress
        activeStress_from_myosin = p['sig_a_N']* tanh(1*uM)  # Nonlinear myosin activation
        # Convert to tensor form (isotropic active stress)
        activeStress_tensor = activeStress_from_myosin*as_tensor([[1, 0], [0, 1]])

        # -------------------------------------------------------------------------------------
        # Constitutive relations for viscoelastic material
        # -------------------------------------------------------------------------------------
        # Total Cauchy stress including active and passive components
        CS = (activeStress_tensor +                                    # Active stress
              p['lmbdaE_N'] * tr(eps) * Identity(2) +                 # Elastic volumetric
              2*p['muE_N']*eps +                                      # Elastic deviatoric
              p['tauc']*p['lmbdaE_N']*(trace-trace_n)/dt*Identity(2) + # Viscous volumetric
              2*p['muE_N']*p['tauc']*(eps-eps_n)/dt)                  # Viscous deviatoric

        # Passive component of Cauchy stress
        CS_passive = (p['lmbdaE_N'] * tr(eps) * Identity(2) +         # Elastic volumetric
                     2*p['muE_N']*eps +                               # Elastic deviatoric
                     p['tauc']*p['lmbdaE_N']*(trace-trace_n)/dt*Identity(2) + # Viscous volumetric
                     2*p['muE_N']*p['tauc']*(eps-eps_n)/dt)           # Viscous deviatoric

        # Calculate positive and negative parts for mechanochemical feedback
        trace_positive = conditional(gt(trace,0),trace,0)*non_opto_cell    # Positive strain
        trace_negative = conditional(gt(0,trace),trace,0)*non_opto_cell    # Negative strain
        J_positive = conditional(gt(J-1,0),J-1,0)*non_opto_cell           # Positive volume change
        detCS_positive = conditional(gt(0,det(CS_passive)),det(CS_passive),0)  # Positive stress determinant

        # Calculate traction force
        tF = kN*u                            # Linear spring force
        tF_mag = kN*sqrt(inner(u,u))        # Magnitude of traction force
        tF_mag_projected = project(tF_mag, K_DG0)  # Project for visualization

        # -------------------------------------------------------------------------------------
        # Handle photoactivation event
        # -------------------------------------------------------------------------------------
        if t == t_opto:
            # Initialize photoactivation pattern
            G_0 = Constant(p['alpha'])*SquareCompartmentDoublet(1,5e-6/p['length_scale'])

            # Set and project new initial conditions after photoactivation
            uG_n = project(G_0, V.sub(0).collapse())
            uR_n = project(R_0, V.sub(1).collapse())
            uM_n = project(M_0, V.sub(2).collapse())
            u_n = project(u0, V.sub(3).collapse())
```

```
            # Update solution vector
            assign(U_tot_n, [uG_n,uR_n,uM_n,u_n])
            uG_n, uR_n, uM_n, u_n = split(U_tot_n)

        # ------------------------------------------------------------------------
        # Define mechanochemical feedback
        # ------------------------------------------------------------------------
        # Check cellularisation parameter
        cellularisation = int(p['cellularisation'])

        # Calculate feedback based on positive strain in non-activated cells
        if cellularisation:
            feedback = p['fb']*trace_positive*non_opto_cell
        else:
            feedback = Constant(0)

        # Disable feedback if feedback strength is small
        if p['fb'] < 0.01:
            feedback = Constant(0)

        # ------------------------------------------------------------------------
        # Define reaction kinetics for signaling cascade
        # ------------------------------------------------------------------------
        # GEF activation/inactivation with mechanical feedback
        React_G = -p['lambda_decay']*uG + feedback
        # RhoA activation by GEF
        React_R = p['b']*(uG - uR)
        # Myosin activation by RhoA
        React_M = p['s']*(uR - uM)

        # ------------------------------------------------------------------------
        # Construct weak form of the coupled system
        # ------------------------------------------------------------------------
        # Reaction-diffusion equations with DG formulation
        FcG = DGWeakFormRD(uG,uG_n,vG,p['DG_N'],u,J,J_n,F,n,dx,dS,dSM,dt,cellularisation) -J*(React_G)
    *vG*dx
        FcR = DGWeakFormRD(uR,uR_n,vR,p['DR_N'],u,J,J_n,F,n,dx,dS,dSM,dt,cellularisation) -J*(React_R)
    *vR*dx
        FcM = DGWeakFormRD(uM,uM_n,vM,p['DM_N'],u,J,J_n,F,n,dx,dS,dSM,dt,cellularisation) -J*(React_M)
    *vM*dx

        # Mechanical equilibrium equation
        F_u = inner(CS,grad(vu))*dx + kN*inner(u,vu)*dx

        # Complete weak form
        FWF = FcG + FcR + FcM + F_u

        # ------------------------------------------------------------------------
        # Configure and solve the nonlinear system
        # ------------------------------------------------------------------------
        # Set solver parameters for SNES (nonlinear solver)
        snes_solver_parameters = {
            "nonlinear_solver": "snes",
            "snes_solver": {
                "linear_solver": "lu",        # Direct LU solver for linear system
                'absolute_tolerance': 1e-6,    # Convergence criteria
                'relative_tolerance': 1e-6,
                "maximum_iterations": 20,      # Limit iteration count
                "report": True,               # Print convergence info
                "error_on_nonconvergence": True
            }
        }

        # Set up nonlinear variational problem
        dFWF = derivative(FWF, U_tot, dU)   # Calculate Jacobian
```

```
        problem = NonlinearVariationalProblem(FWF, U_tot, [], J=dFWF)
        solver = NonlinearVariationalSolver(problem)
        solver.parameters.update(snes_solver_parameters)
        info(solver.parameters, False)

        # Solve the system
        (iter, converged) = solver.solve()




        # -------------------------------------------------------------------------------------
        # Prepare and save solution fields for visualization
        # -------------------------------------------------------------------------------------
        # Copy current solutions for post-processing
        U_tot_save.assign(U_tot)                # Current solution
        U_tot_save_n.assign(U_tot_n)            # Previous solution
        _uG, _uR, _uM, _u = U_tot_save.split(deepcopy=True)  # Split into components

        # -------------------------------------------------------------------------------------
        # Project solution fields onto appropriate function spaces
        # -------------------------------------------------------------------------------------
        # Vector fields (CG1 space)
        disp.assign(project(_u, V_CG1))              # Displacement field
        TractionF.assign(project(tF, V_CG1))         # Traction force

        # Molecular species concentrations (DG0 space)
        GEF.assign(project(_uG, K_DG0))              # GEF concentration
        RhoA.assign(project(_uR, K_DG0))             # RhoA concentration
        Myosin.assign(project(_uM, K_DG0))           # Myosin concentration

        # Domain markers and geometric quantities (DG0 space)
        activatedCell.assign(project(cell1, K_DG0))           # Activated cell region
        nonOptoCells.assign(project(non_opto_cell, K_DG0))    # Non-photoactivated regions
        pattern.assign(project(kN, K_DG0))                    # Micropattern
        Jacobian.assign(project(J, K_DG0))                    # Volume change
        JacobianPositive.assign(project(J_positive, K_DG0))   # Positive volume change
        feedbackPositive.assign(project(feedback, K_DG0))     # Mechanical feedback

        # Strain measures (DG0 space)
        traceGreenLagrange.assign(project(trace, K_DG0))                        # Volumetric strain
        traceGreenLagrangePositive.assign(project(trace_positive, K_DG0))      # Positive strain
        detCauchyStressPositive.assign(project(detCS_positive, K_DG0))         # Positive stress

        # Stress and strain tensors (DG0 tensor space)
        Cauchystress.assign(project(CS, W_DG0))                     # Total Cauchy stress
        Cauchystress_passive.assign(project(CS_passive, W_DG0))     # Passive stress
        activeStress.assign(project(activeStress_tensor, W_DG0))    # Active stress
        strainGreenLagrange.assign(project(eps, W_DG0))             # Strain tensor
        defGrad_save.assign(project(F, W_DG0))                      # Deformation gradient
        CauchyGreenInverse_save.assign(project(C1, W_DG0))          # Inverse Cauchy-Green tensor

        # -------------------------------------------------------------------------------------
        # Write fields to XDMF file for visualization
        # -------------------------------------------------------------------------------------
        # Vector fields
        xdmf_file.write(disp, t)                 # Displacement
        xdmf_file.write(TractionF, t)            # Traction

        # Molecular concentrations
        xdmf_file.write(GEF, t)                  # GEF
        xdmf_file.write(RhoA, t)                 # RhoA
        xdmf_file.write(Myosin, t)               # Myosin

        # Domain markers and geometric quantities
        xdmf_file.write(activatedCell, t)        # Activated cells
```

```
        xdmf_file.write(nonOptoCells, t)          # Non-photoactivated cells
        xdmf_file.write(pattern, t)               # Micropattern
        xdmf_file.write(Jacobian, t)              # Volume change
        xdmf_file.write(JacobianPositive, t)      # Positive volume change
        xdmf_file.write(feedbackPositive, t)      # Mechanical feedback

        # Strain measures
        xdmf_file.write(traceGreenLagrange, t)             # Total strain
        xdmf_file.write(traceGreenLagrangePositive, t)     # Positive strain
        xdmf_file.write(detCauchyStressPositive, t)        # Positive stress

        # Stress and strain tensors
        xdmf_file.write(Cauchystress, t)                    # Total stress
        xdmf_file.write(Cauchystress_passive, t)            # Passive stress
        xdmf_file.write(activeStress, t)                    # Active stress
        xdmf_file.write(strainGreenLagrange, t)             # Strain tensor
        xdmf_file.write(defGrad_save, t)                    # Deformation gradient
        xdmf_file.write(CauchyGreenInverse_save, t)         # Inverse Cauchy-Green

        # ---------------------------------------------------------------------------------------
        # Calculate and store derived quantities
        # ---------------------------------------------------------------------------------------
        # Store current time
        time_array.append(t)

        # Calculate strain energy in different regions
        dx1 = Measure('dx', domain=mesh, subdomain_data=subdomains, subdomain_id=1)
        strainEnergyLeft = calculateStrainEnergy(u, kN, dx1)      # Left cell

        dx2 = Measure('dx', domain=mesh, subdomain_data=subdomains, subdomain_id=2)
        strainEnergyRight = calculateStrainEnergy(u, kN, dx2)     # Right cell

        strainEnergyTotal = calculateStrainEnergy(u, kN, dx)      # Total system

        # Extract boundary curve for shape analysis
        boundary_curve = get_boundary_of_deformed_mesh(_u, geo_file_name)

        # Store energy and boundary data
        strainEnergyLeft_vs_time.append(strainEnergyLeft)
        strainEnergyRight_vs_time.append(strainEnergyRight)
        strainEnergyTotal_vs_time.append(strainEnergyTotal)
        boundary_curve_vs_time.append(boundary_curve)

        # ---------------------------------------------------------------------------------------
        # Update time step and solution
        # ---------------------------------------------------------------------------------------
        t = t + DT                     # Increment time
        U_tot_n.assign(U_tot)          # Store current solution for next step

        # ---------------------------------------------------------------------------------------
        # Save simulation data to files
        # ---------------------------------------------------------------------------------------
        # Create dictionary of energy data
        outputDict = {
            'time': np.array(time_array),
            'strainEnergyLeft': np.array(strainEnergyLeft_vs_time),
            'strainEnergyRight': np.array(strainEnergyRight_vs_time),
            'strainEnergyTotal': np.array(strainEnergyTotal_vs_time)
        }

        # Save energy data to CSV
        pd.DataFrame.from_dict(data=outputDict).to_csv(path_to_rawData+'strainEnergy.csv', header=True)

        # Save boundary curve data to NPZ file
        np.savez(path_to_rawData + 'boundary_curve.npz',
```

```
                 boundary_curve=boundary_curve_vs_time,
                 time=np.array(time_array))

# Execute simulation if script is run directly
if __name__ == "__main__":
    simulation()
```
